# Structures of wild-type and a constitutively closed mutant of connexin26 shed light on channel regulation by CO_2_

**DOI:** 10.1101/2023.08.22.554292

**Authors:** Deborah H. Brotherton, Sarbjit Nijjar, Christos G. Savva, Nicholas Dale, Alexander D. Cameron

## Abstract

Connexins allow intercellular communication by forming gap junction channels (GJCs) between juxtaposed cells. Connexin26 (Cx26) can be regulated directly by CO_2_. This is proposed to be mediated through carbamylation of K125. We show that mutating K125 to glutamate, mimicking the negative charge of carbamylation, causes Cx26 GJCs to be constitutively closed. Through cryo-EM we observe that the K125E mutation pushes a conformational equilibrium towards the channel having a constricted pore entrance, similar to effects seen on raising the partial pressure of CO_2_. In previous structures of connexins, the cytoplasmic loop, important in regulation and where K125 is located, is disordered. Through further cryo-EM studies we trap distinct states of Cx26 and observe density for the cytoplasmic loop. The interplay between the position of this loop, the conformations of the transmembrane helices and the position of the N-terminal helix, which controls the aperture to the pore, provides a mechanism for regulation.

## Introduction

Connexins form hexameric channels in the plasma membrane known as hemichannels, which can either function as regulated passageways between the cell and its environment, or dock with a hemichannel from another cell to form a dodecameric intercellular channel, or gap junction channel (GJC). Connexins have been shown to be directly regulated by various stimuli such as voltage (Valiunas, 2002; Young & Peracchia, 2004), pH (Bevans & Harris, 1999; Khan et al., 2020; Yu et al., 2007) or indirectly via intracellular calcium ion concentrations (Peracchia, 2004). Recent reports based on structural data also suggest that lipids may be involved in regulation (H. J. Lee et al., 2023; S. N. Lee et al., 2023; Qi, Acosta Gutierrez, et al., 2023). We have shown, however, that connexin26 (Cx26) and other similar β-connexins (Cx30, Cx32), can be regulated by the direct action of physiological concentrations of carbon dioxide independently of pH (Huckstepp et al., 2010; Meigh et al., 2013). Mutants of Cx26 are a leading cause of congenital deafness (Xu & Nicholson, 2013). While many of the mutations are non-syndromic, others lead to severe diseases such as keratitis ichthyosis deafness syndrome (KIDS) (Xu & Nicholson, 2013). Hemichannels are known to have different properties to GJCs (Stout et al., 2004) and our previous results show that an increase in the partial pressure of CO_2_ (PCO_2_) will open Cx26 hemichannels (Huckstepp et al., 2010) but close Cx26 GJCs (Nijjar et al., 2021), which are open under physiological conditions of PCO_2_.

There are 20 connexin genes in the human genome (Abascal & Zardoya, 2013) and several structures have now been published (Bennett et al., 2016; Brotherton et al., 2022; Flores et al., 2020; Khan et al., 2020; H. J. Lee et al., 2023; Lee et al., 2020; S. N. Lee et al., 2023; Maeda et al., 2009; Myers et al., 2018; Qi, Acosta Gutierrez, et al., 2023; Qi, Lavriha, et al., 2023). The connexin subunit, which is common to all, consists of four transmembrane helices (TMs) with a cytoplasmic N-terminal helix that in the hexameric arrangement of the hemichannel points towards the central pore (Maeda et al., 2009). In structures of the dodecameric GJC, the extracellular part, involved in docking is well defined, whereas the cytoplasmic region is much more variable. A large cytoplasmic loop between TM2 and TM3, shown to be involved in regulation, has not been visible in any structure. Structures of the connexins either have the N-terminal helices tucked back against the wall of the channel (Flores et al., 2020; H. J. Lee et al., 2023; S. N. Lee et al., 2023; Myers et al., 2018), in a raised position (H. J. Lee et al.; Lee et al., 2020; Qi, Acosta Gutierrez, et al., 2023), in an intermediate position (Brotherton et al., 2022; H. J. Lee et al., 2023; Maeda et al., 2009) or not well-defined in the density (Bennett et al., 2016; Brotherton et al., 2022; Khan et al., 2020; S. N. Lee et al., 2023; Qi, Lavriha, et al., 2023). The position of the N-terminal helix is thought to be important in the regulation of channel permeability. We have shown previously for human Cx26 GJCs, that the position of the N-terminus is dependent on the partial pressure of CO_2_ (PCO_2_) at constant pH (Brotherton et al., 2022). By examining structures from protein vitrified at different levels of PCO_2_, we observed that under conditions of high PCO_2_ the conformation of the protein was biased towards a conformation where the N-terminus protrudes radially into the pore to form a constriction at the centre (N_Const_). Two distinct conformations of the protein were seen with the predominant difference between them in the cytoplasmic portion of TM2 (Brotherton et al., 2022) where an anticlockwise rotation of TM2 correlated with more definition of the density for the N-terminus. On the other hand, under low PCO_2_ conditions the conformation with the more defined N-terminus was not observed and the channel appeared more open (N_Flex_).

Based on a wealth of mutational data it has been hypothesised that the regulation of hemichannel opening by CO_2_ is through a carbamylation reaction of a specific lysine (Meigh et al., 2015; Meigh et al., 2013; Nijjar et al., 2021). This post-translational modification is a reversible and highly labile reaction of CO_2_ (Lorimer, 1983) that effectively changes the charge of a neutral lysine residue to make it negative. A so-called “carbamylation motif” was identified in CO_2_-sensitive connexins (Dospinescu et al., 2019; Meigh et al., 2013) that when introduced into a related CO_2_-insensitive connexin rendered the protein CO_2_-sensitive (Meigh et al., 2013). In Cx26 this motif has the sequence K_125_VRIEG_130_. In the crystal structure of the Cx26 GJC that was published in 2009 (Maeda et al., 2009), Lys125, which is conserved amongst β-connexins that are known to be modulated by CO_2_ (Dospinescu et al., 2019), is positioned near to the N-terminus of TM3 within ∼6Å of Arg104 of TM2 of the neighbouring subunit, at either side of the disordered cytoplasmic loop. It was suggested that upon carbamylation, the negative charge of the modified lysine would attract Arg104 causing a conformational change (Meigh et al., 2013). In hemichannels, mutation of Lys125 to glutamate, so mimicking the charge of the carbamylated lysine (K125E) results in constitutively open hemichannels consistent with elevated PCO_2_, whereas the corresponding K125R mutation results in hemichannels that cannot be opened by CO_2_ (Meigh et al., 2013). In GJCs, the K125R mutation results in the protein not closing in response to CO_2_, though importantly, this mutation does not prevent closure by acidification (Nijjar et al., 2021).

While our previous structures (Brotherton et al., 2022) demonstrated an effect of PCO_2_ on the conformation of the protein, neither Lys125 nor Arg104 were visible in the density. Here we probe this further. By solubilising the protein in the detergent lauryl maltose neopentyl glycol (LMNG) rather than dodecyl β-D-maltoside (DDM) we obtain much improved density for the cytoplasmic region of the protein. We refine two conformationally diverse structures of the protein, where we see much more defined differences in the cytoplasmic region of the protein than we were able to observe previously. These data suggest a mechanism for closure involving the concerted movements of TM2 and the KVRIEG motif. We show that the structure with the K125E mutation matches the more closed conformation of the protein.

## Results

### Conformations of TMs 1, 2 and the KVRIEG motif correlate with position of the N-terminus

In an attempt to improve the resolution of the cytoplasmic region of the protein we changed the method of solubilisation and purification by substituting DDM with LMNG and swapping from phosphate buffers to CO_2_/HCO_3_^-^ buffers throughout the process. The switch from DDM to LMNG was intended to provide clarity on the provenance of density within the pore of the protein that was observed in our previous structures (Brotherton et al., 2022) and which was considered to be either DDM, as an artefact of solubilisation, or lipid. Lipids have been observed in the pore of other connexins and related large-pored channels and have been suggested to be part of the mechanism (Burendei et al., 2020; H. J. Lee et al., 2023; Lee et al., 2020; S. N. Lee et al., 2023). Our use of high CO_2_/HCO_3_^-^ buffers throughout purification was intended to keep the gap junction in the closed state throughout, and hence reduce the chance of extraneous lipid or detergent entering the channel pore.

Data were collected from Cx26 vitrified at a PCO_2_ corresponding to 90mmHg. Refinement with D6 symmetry imposed resulted in a map with a nominal resolution of 2.0Å as defined by gold standard Fourier Shell Correlations (Rosenthal & Henderson, 2003; Scheres, 2012) (Figure 1 – figure supplement 1, Table 1). This was further classified using the procedure that we previously developed, involving particle expansion and signal subtraction, to focus on just the cytoplasmic region of one of the two docked hemichannels (Figure 1 – figure supplements 1 and 2 (Brotherton et al., 2022)).

**Figure 1:**
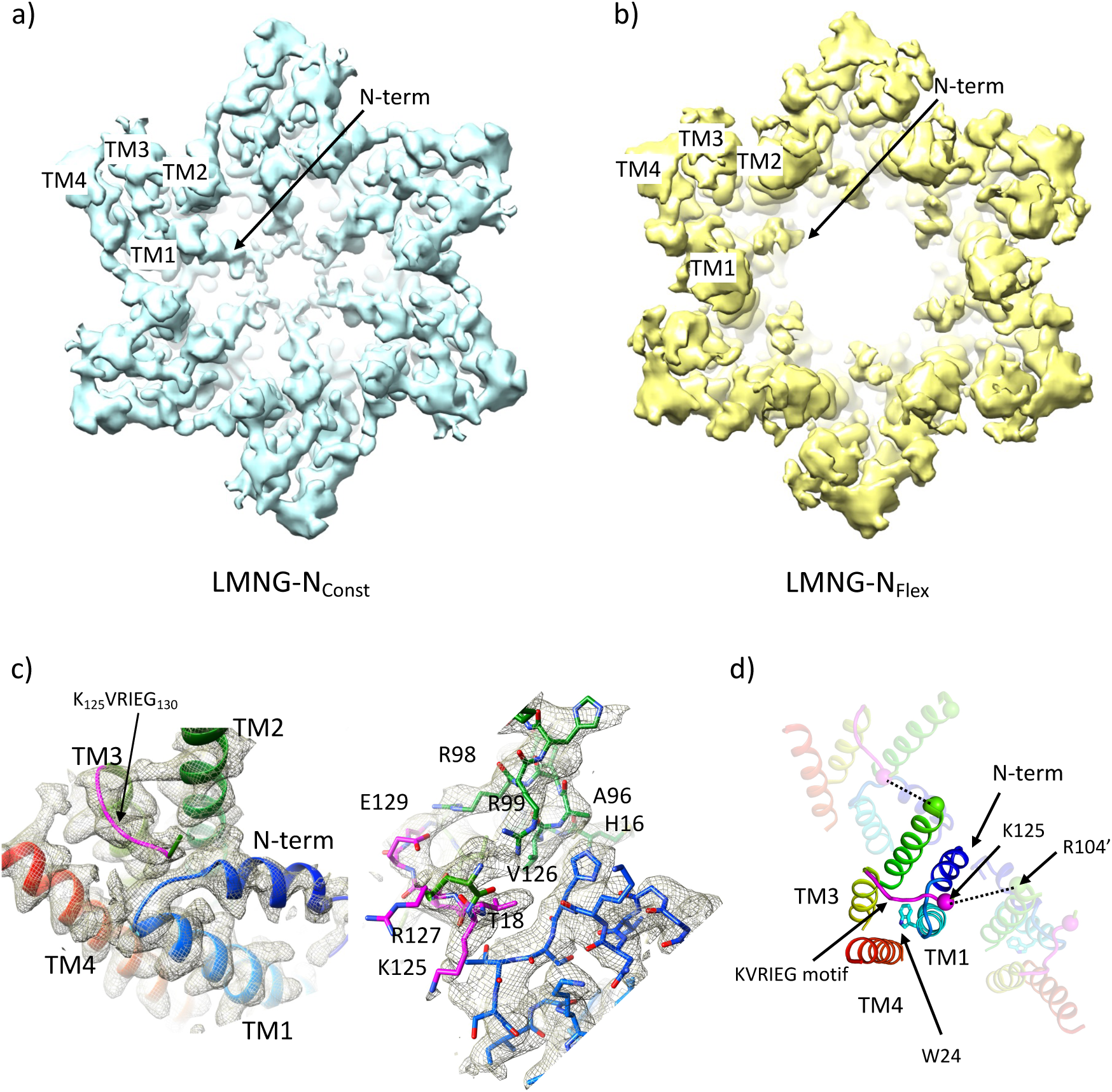
Distinct classes from classification of Cx26 solubilised in LMNG. **a)** Overall density associated with LMNG-N_Const_ viewed from the cytoplasmic face. **b)** Density associated with LMNG-N_Flex_. **c)** As (a) focussed on the KVRIEG motif and the link between the N-terminus and TM1. Left: the cartoon has been coloured through the colours of the rainbow with blue at the N-terminus to red at the C-terminus, except for the KVRIEG motif, which is shown in magenta. Right: stick representation with the same colouring showing the interactions between the residues on the link between the N-terminus and TM1 (blue), residues on TM2 (green) and the KVRIEG motif (magenta). **d)** Cartoon representation of the cytoplasmic region of the LMNG-N_Const_ structure. The two neighbouring subunits to the central subunit in the figure have been made semi-transparent. The dotted lines show the proximity of K125 of one subunit to R104 of the neighbouring subunit. Trp24 on TM1 is in the region of TM1 that adopts an altered conformation with respect to the previously solved structures of Cx26.

**Table 1:** Cryo-EM data collection and processing statistics.

|  | **K125E<sub>90</sub>** | **K125R<sub>90</sub>** | **LMNG<sub>90</sub>** | **K125E<sub>HEPES</sub>** | **WT<sub>HEPES</sub>** |
| --- | --- | --- | --- | --- | --- |
| Voltage (kV) | 300 | 300 | 300 | 300 | 300 |
| Magnification (x1000) X | 105 | 105 | 105 | 75 | 75 |
| Camera | K3 | K3 | K3 | Falcon 3 | Falcon 3 |
| Frame alignment on Falcon 3 | - |  | - | yes | yes |
| Camera mode | Super-Resolution | Counting bin1 | Super-Resolution bin2 | counting | counting |
| Energy filter (eV) | 20 | 20 | 20 | 20 | 20 |
| Defocus range (μm) | -0.8 to -2.0 | -0.8 to -2.0 | -0.8 to -2.3 | -0.3 to -1.7 | -0.5 with Volta phaseplate |
| Pixel size (Å/pix) | 0.85 | 0.835 | 0.835 | 1.08 | 1.08 |
| Dose on detector (e<sup>-</sup>/pix/sec) | 10 | 15 | 18.77 |  | 0.7 |
| Dose on sample (e<sup>-</sup>/pix/sec) | 11.4 | 18 |  | 1.06 | 0.69 |
| Exposure time (sec) | 3 | 2 | 2 | 44.01 | 60 |
| No. of images | 4731 | 10044 | 11362 | 2573 | 2436 |
| Frames per image | 45 | 50 | 50 | 40 | 75 |
| Final particle number | 189887 | 222622 | 204438 | 147546 | 60995 |
| **Resolution<sup>a</sup>** |  |  |  |  |  |
| masked D6 (Å) | 2.2 | 2.1 | 2.0 | 4.3 | 4.9 |

<sup>a)</sup> From Relion\_postprocess (Scheres, 2012)

The results from this classification were broadly in line with our previous results. However improved definition of the density in the cytoplasmic region enabled us to model this region more accurately. As before, the position of the cytoplasmic region of TM2 in these maps correlated with the presence or absence of the N-terminus. Maps from two classifications based on the most extreme positions of TM2 and corresponding clarity of the N-terminus were taken forward for further analysis. (Figure 1 – figure supplement 1, Table 2). These maps, both of which have a similar resolution of 2.3Å, have been respectively denoted LMNG-N_Const_ (N-terminus defined and constricting the pore) and LMNG-N_Flex_ (N-terminus not visible) following the nomenclature above (Figure 1). As observed in previous maps, density associated with a hydrophobic molecule was present in the pore of both maps (Figure 1 – figure supplement 3a).

**Table 2.** Cryo-EM refinement and validation statistics.

|  | K125E<sub>90</sub> | LMNG<sub>90</sub> |  |  |  |
| --- | --- | --- | --- | --- | --- |
|  |  | N<sub>Flex</sub> | N<sub>Const</sub> | N<sub>Const-mon</sub> | N<sub>Flex-mon</sub> |
| Deposited Structure PDB ID | 8Q9Z | 8QA1 | 8QA0 | 8QA2 | 8QA3 |
| Final particle number | 161625 | 59005 | 35007 | 357859 | 240137 |
| **Map resolution** |  |  |  |  |  |
| FSC threshold | 0.143 | 0.143 | 0.143 | 0.143 | 0.143 |
| Symmetry imposed<sup>a</sup> | C6 | C6 | C6 | C1 | C1 |
| Unmasked (Å) | 2.3 | 2.2 | 2.4 | 2.6 | 2.4 |
| Masked (Å) (masked) | 2.4 | 2.0 | 2.1 | 2.3 | 2.2 |
| **Refinement** |  |  |  |  |  |
| Initial model (PDB code) | 7QEQ | 7QEQ | K125E<sub>90</sub> | AlphaFold | mon-PCN |
| Resolution (Å; FSC=0.5) Model | 2.6 | 2.4 | 2.6 | 2.6 | 2.6 |
| Sharpening B factor (Å<sup>2</sup>) | local | local | local | local | local |
| **Model composition** | 12 chains | 12 chains | 12 chains | 12 chains | 12 chains |
| Non-hydrogen atoms | 19704 | 18720 | 19878 | 18945 | 18580 |
| Protein residues | 2316 | 2190 | 2340 | 2260 | 2226 |
| Water | 348 | 318 | 342 | 322 | 254 |
| Ligand: lipid/detergent | 48 | 24 | 24 | 13 | 12 |
| **B factor (Å<sup>2</sup>)** |  |  |  |  |  |
| protein | 66 | 54 | 77 | 68 | 73 |
| water | 42 | 33 | 56 | 45 | 44 |
| Lipid/detergent | 56 | 60 | 85 | 69 | 70 |
| **R.m.s. deviations** |  |  |  |  |  |
| Bond lengths (Å) | 0.004 | 0.002 | 0.004 | 0.002 | 0.003 |
| Bond angles (°) | 0.516 | 0.484 | 0.593 | 0.450 | 0.544 |
| **Validation** |  |  |  |  |  |
| MolProbity score | 1.54 | 1.25 | 1.55 | 1.53 | 1.74 |
| Clashscore | 5.66 | 3.46 | 6.50 | 4.90 | 5.20 |
| Rotamer outliers (%) | 1.83 | 1.42 | 1.05 | 2.32 | 3.71 |
| **Ramachandran plot** |  |  |  |  |  |
| Favoured (%) | 97.84 | 98.97 | 96.99 | 98.5 | 97.89 |
| Allowed (%) | 2.16 | 1.03 | 3.01 | 1.49 | 2.11 |
| Disallowed (%) | 0 | 0 | 0 | 0 | 0 |
| **CaBlam Outliers (%)** | 1.53 | 1.24 | 1.6 | 0.83 | 1.08 |
| **Correlation coefficients** |  |  |  |  |  |
| CC (mask) | 0.86 | 0.87 | 0.85 | 0.86 | 0.87 |
| CC (box) | 0.7 | 0.71 | 0.66 | 0.68 | 0.70 |
| CC (peaks) | 0.68 | 0.70 | 0.63 | 0.66 | 0.68 |
| CC (volume) | 0.84 | 0.85 | 0.84 | 0.85 | 0.85 |
| Mean CC for ligands | 0.71 | 0.73 | 0.61 | 0.68 | 0.70 |

For the LMNG-N_Const_ conformation, the density for the side-chains of the residues of the N-terminus and the following link to TM1 is much clearer than seen in the other maps (Figure 1c), however it remains difficult to place the first three residues of the N-terminus unambiguously. This new structure is an advance on the previous structure of the equivalent conformation obtained in DDM (PDB 7QEW) (Figure 1 – figure supplement 3c). In addition to being able to assign more residues to the density in LMNG-N_Const_, there are two main regions which have been modelled differently. The first area that differs is in TM1 (Figure 1d, Figure 1 – figure supplement 3b). Previously we noted variation between the Cx26 crystal structures and our cryo-EM structures in the position of the residues between Val37 and Glu42 (Brotherton et al., 2022). In the LMNG-N_Const_ structure, the N-terminal region of TM1, preceding this area and comprising residues Gly21 to Phe31, is rotated with respect to that modelled previously, changing the position of the π-helix in TM1 from residues Ile20-Leu25 to residues Phe29-Val38 (Figure 1 – figure supplement 3b). Thus, the conformation of TM1 in the LMNG-N_Const_ structure differs from the crystal structures at both the N-terminal and C-terminal ends through variations in the twist of the helical repeats.

The second difference is at the cytoplasmic side of TM3. In all structures of connexins solved to date the cytoplasmic loop has been disordered. For Cx26 this region extends from approximately residue Arg104 at the C-terminus of TM2 to Glu129 at the N-terminus of TM3. In the crystal structure (Cx26-xtal) (Maeda et al., 2009) residues 125 to 129 have been modelled as part of TM3, whereas in our previous structures solved by cryo-EM (Brotherton et al., 2022) there was no evidence in the associated maps of the helix extending beyond Glu129. Instead, when analysing these maps it was noted that unassigned density protruded from near to the top of TM3 towards the loop between the N-terminus and TM1 in a manner that resembles models from AlphaFold (Jumper et al., 2021; Varadi et al., 2022). In the LMNG-N_Const_ map this density is much more clearly defined as the C-terminal end of the cytoplasmic loop as it joins onto TM3 (Figure 1c, Figure 1 – figure supplement 2). Residues Gln124 to Ile128, which form part of the K_125_VRIEG_130_ motif, important for carbamylation, were modelled into it, with Val126 located just above the linking region between the N-terminal helix and TM1 (Figure 1c). Though the density for the side chains is poor and there is no definitive interaction, Lys125 in this position is relatively close to the side chain of Arg104 of the neighbouring subunit, with which it had been proposed to form a salt bridge following carbamylation (Figure 1d).

By contrast, in the LMNG-N_Flex_ map neither the N-terminus nor the KVRIEG motif are well defined (Figure 1b). The LMNG-N_Flex_ map is reminiscent of the map derived from Cx26 particles vitrified under low PCO_2_ conditions (Brotherton et al., 2022). In the associated structure the conformation of TM1 is the same as was modelled for the previous structures in DDM. A comparison between the two conformations derived from the new data is shown in Figure 2a and 2b and Figure 2 – videos 1 and 2. The conformational change in TM1 results in the side-chain of Trp24 rotating by ∼90° between the two extreme positions (Figure 2b, Figure 2 – video 2). At one of these extremes, it faces the exterior of the protein, and nestles a detergent or lipid tail. At the other extreme, it is within the core of the protein next to Arg143 and Ala88 (Figure 2b). The conformation in the N_Flex_ structure would not be compatible with the position adopted by TM2 in the LMNG-N_Const_ structure because Thr86 and Leu89 would clash with Phe31 and Ile30 on TM1 of the neighbouring subunit (Figure 2b, Figure 2 – videos 1 and 2). The rotation of TM1 changes not only how TM1 interacts with the N-terminus, but also the conformation of the linker between the N-terminal helix and TM1 (Figure 2a). This in turn would not be compatible with the conformation of the KVRIEG motif in LMNG-N_Const_ (Figure 2b). Overall, therefore the constriction of the channel by the N-terminal helix is associated with changes in the positions of TMs 1, 2 and the KVRIEG motif of the cytoplasmic loop.

**Figure 2:**
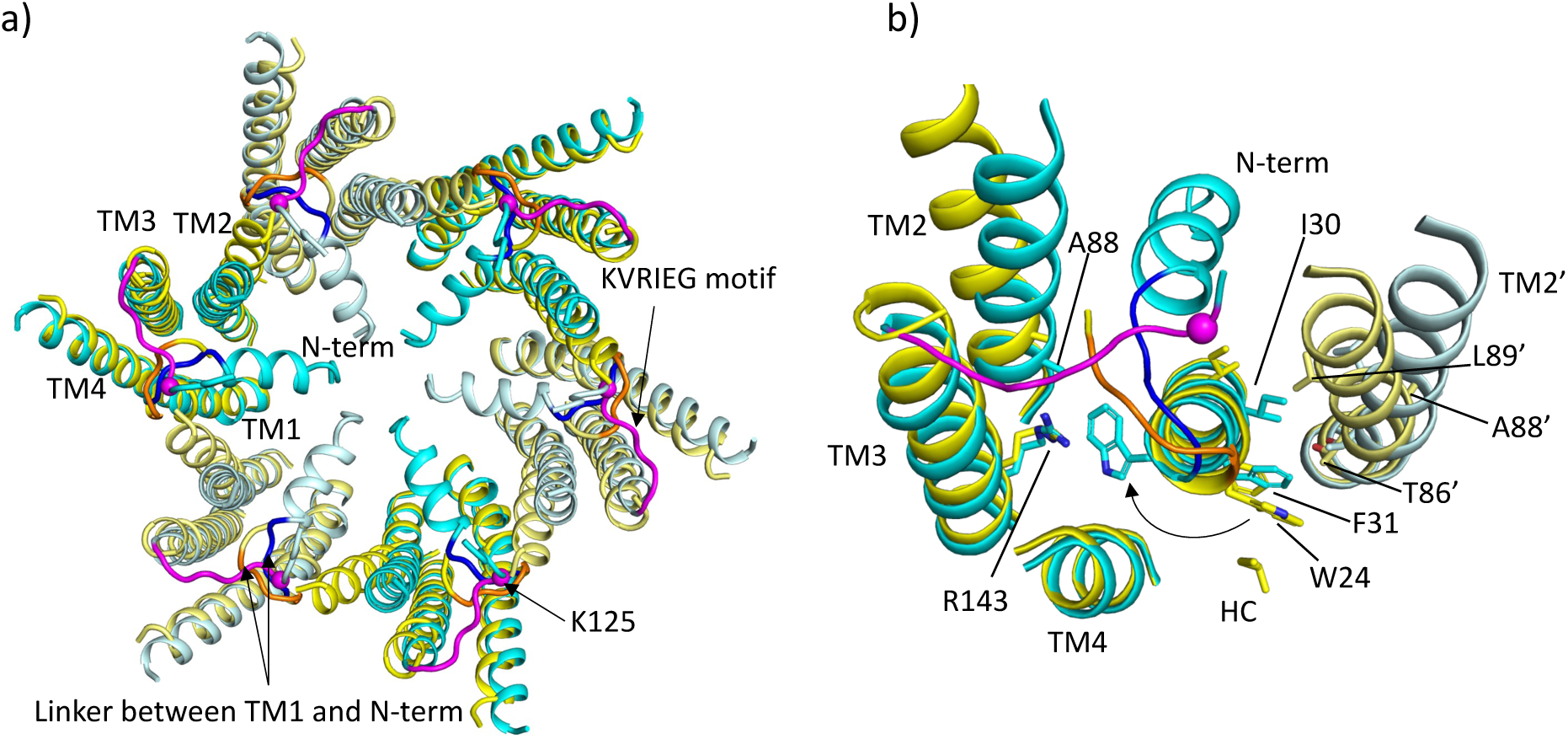
Comparison of LMNG-N_Const_ and LMNG-N_Flex_ structures. **a)** Overall superposition showing the movement of TM2 and the link between the N-terminus and TM1. LMNG-N_Const_ in cyan and LMNG-N_Flex_ in yellow (alternate subunits have been coloured in lighter shades). The KVRIEG motif has been coloured magenta with a sphere indicating the position of K125. The residues between the N-terminus and TM1 for the LMNG-N_Const_ structure have been coloured blue. **b)** As (a) but focussed on TM1. The conformation of TM1 differs between the two structures as seen by the change in position of Trp24. TM2’ is from the neighbouring subunit. HC denotes the hydrocarbon chain from a lipid. The positions of T86’ and L89’ of TM2 in the N_Flex_ conformation are not compatible with F31 and I30 TM1 in the N_Const_ conformation.

### Density for cytoplasmic loop compatible with models from AlphaFold

Despite the LMNG-N_Const_ map being much clearer for the cytoplasmic region of the protein residues from 107 to 123 were still missing. We, therefore, carried out another classification of the particles focussed on the cytoplasmic region of a single subunit (see methods). As above, this resulted in a range of maps showing varying positions of the transmembrane helices and clarity of the N-terminal helix (Figure 3 – figure supplement 1). Importantly, in one case and where the N-terminus was clearly defined, extra density was also seen for the cytoplasmic loop, albeit at low resolution. The structure was tentatively built into the density with the cytoplasmic loop of the classified subunit in a conformation resembling models from AlphaFold (Jumper et al., 2021) (Figure 3) and with a complete N-terminus (Table 2, Figure 3 – figure supplement 2). Only residues 109 to 114 were omitted in the final structure as the placement of these residues was ambiguous. In the maps associated with this structure there is density that we cannot assign, near to Lys125, between Ser19 in the TM1-N-term linker, Tyr212 of TM4 and Tyr97 on TM3 of the neighbouring subunit, which may be a small molecule that has bound. Overall, the conformation of the subunit is very similar to the LMNG-N_Const_ structure from the classification based on the masked hemichannels with an RMSD of 0.38Å for 198 Cα atoms. A second structure was also refined that had a conformation much more similar to the LMNG-N_Flex_ structure (LMNG-N_Flex-mon_) (RMSD 0.45 for 176 Cα atoms) (Table 2 Figure 3 – figure supplements 1 and 2). Rather surprisingly, given the success of the hemichannel-mask based classification, only the subunit upon which the focussed classification had been carried out had this conformation, with the density from the other subunits appearing more like the map before focussed classification. When hexameric symmetry was applied to the subunit, though the conformation of the N-terminus caused the aperture of the pore to appear closed, steric clashes involving the N-terminal residues suggested a symmetric arrangement of this conformation would not be possible (Figure 3d). This analysis suggested a picture of a flexible molecule that can be captured in different conformations ranging from closed to open but with limited cooperativity between the subunits of the hexamer.

**Figure 3:**
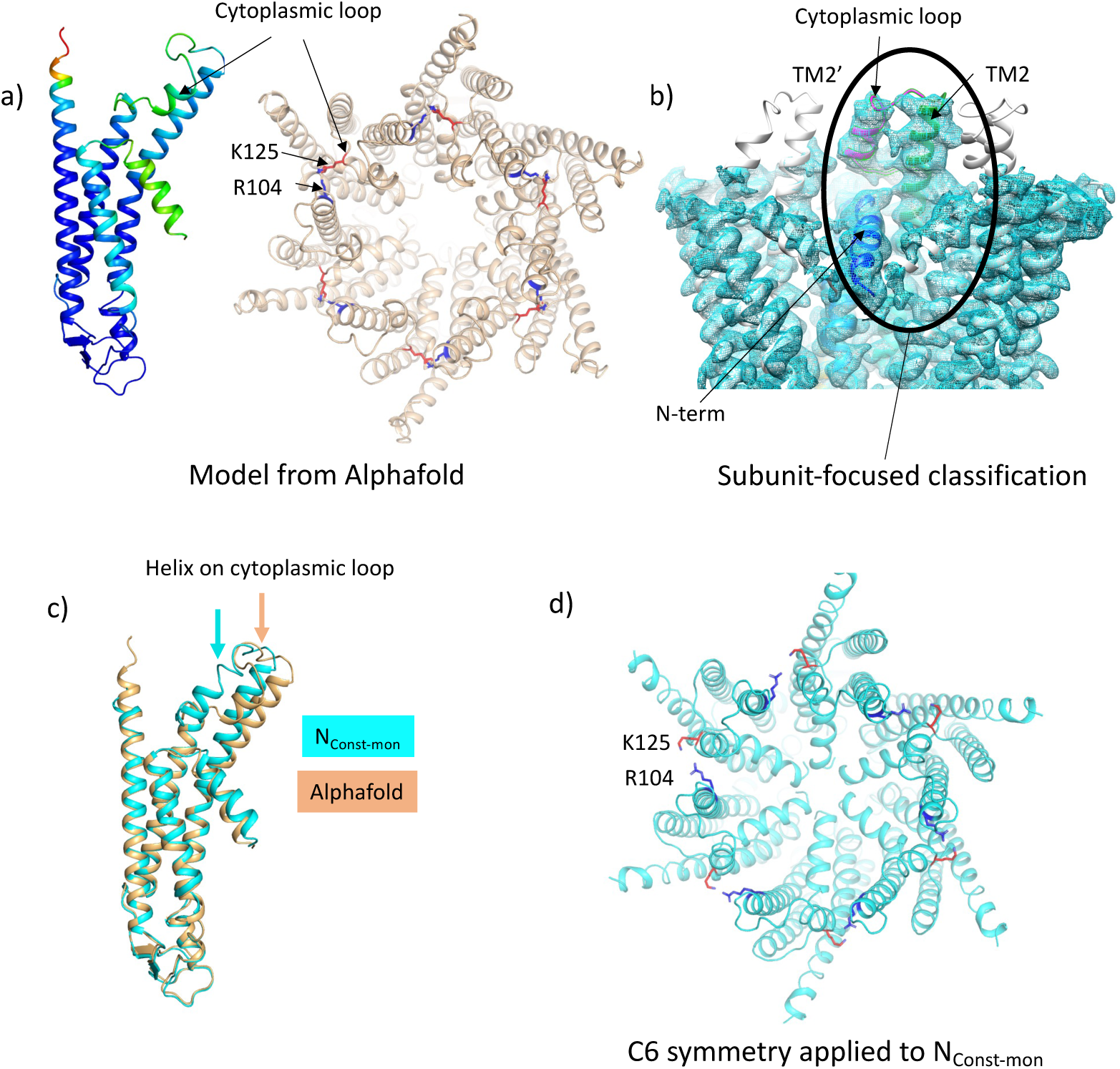
Focussed classification of a single subunit results in density for the cytoplasmic loop consistent with models from AlphaFold. **a)** Models generated by AlphaFold for a single subunit (left; coloured according to confidence level) and for the hexamer (right; in wheat with the position of K125 depicted by red sticks and the position of R104 in blue). **b)** Focussed classification of a single subunit (highlighted by an oval and coloured as in Figure 1d with the cytoplasmic loop in magenta) resulted in clear density for part of the cytoplasmic loop in a conformation consistent with the models from AlphaFold. This does not extend to the neighbouring subunits (coloured grey). **c)** Superposition of the single subunit built into the density (cyan) on the AlphaFold model (wheat). Showing the change in position of the helix in the cytoplasmic loop (highlighted by an arrow in the relevant colour). **d)** Reconstituting a hexamer by replicating the conformation of the subunit seen in (b) to all 6 subunits of the hexamer results in an apparently more closed conformation of the hemichannel, though there are also residue clashes, especially at the N-terminus. Lys 125 and Arg 104 are depicted with red and blue sticks respectively.

### Mutation of K125 to glutamic acid results in constitutively closed GJCs

The above data clearly showed two conformations, from which we could infer a mechanism for closure of the pore. We had previously shown that in GJCs, the K125R mutation remains in the open state even under conditions of high PCO_2_ (Nijjar et al., 2021). Based on our experience with mutations of Cx26 we hypothesised that, if K125E results in constitutively open hemichannels, the same mutation would result in constitutively closed GJCs. Thus, if it were to be true, we could investigate the structure of the proteins under identical buffer conditions where the channel was biased towards open or closed conformations. To verify the effect of mutating Lys125 to a glutamic acid, we used an established dye transfer assay between coupled cells to assess gap junction function (Nijjar et al., 2021). For wild type Cx26, gap junctions readily form between cells and allow rapid transfer of dye from a donor (filled via a patch pipette) to a coupled acceptor cell at a PCO_2_ of 35 mmHg (Figure 4a). Cx26^K125E^ forms structures that resemble wild type gap junctions (Figure 4b). However, these gap junctions appeared to be shut and did not permit dye transfer at a PCO_2_ of 35 mmHg (Figure 4b,d). As the action of an increase in PCO_2_ is to close the wild type Cx26 gap junction, unsurprisingly Cx26^K125E^ gap junctions remained closed at a PCO_2_ of 55 mmHg (Figure 4c,d).

**Figure 4:**
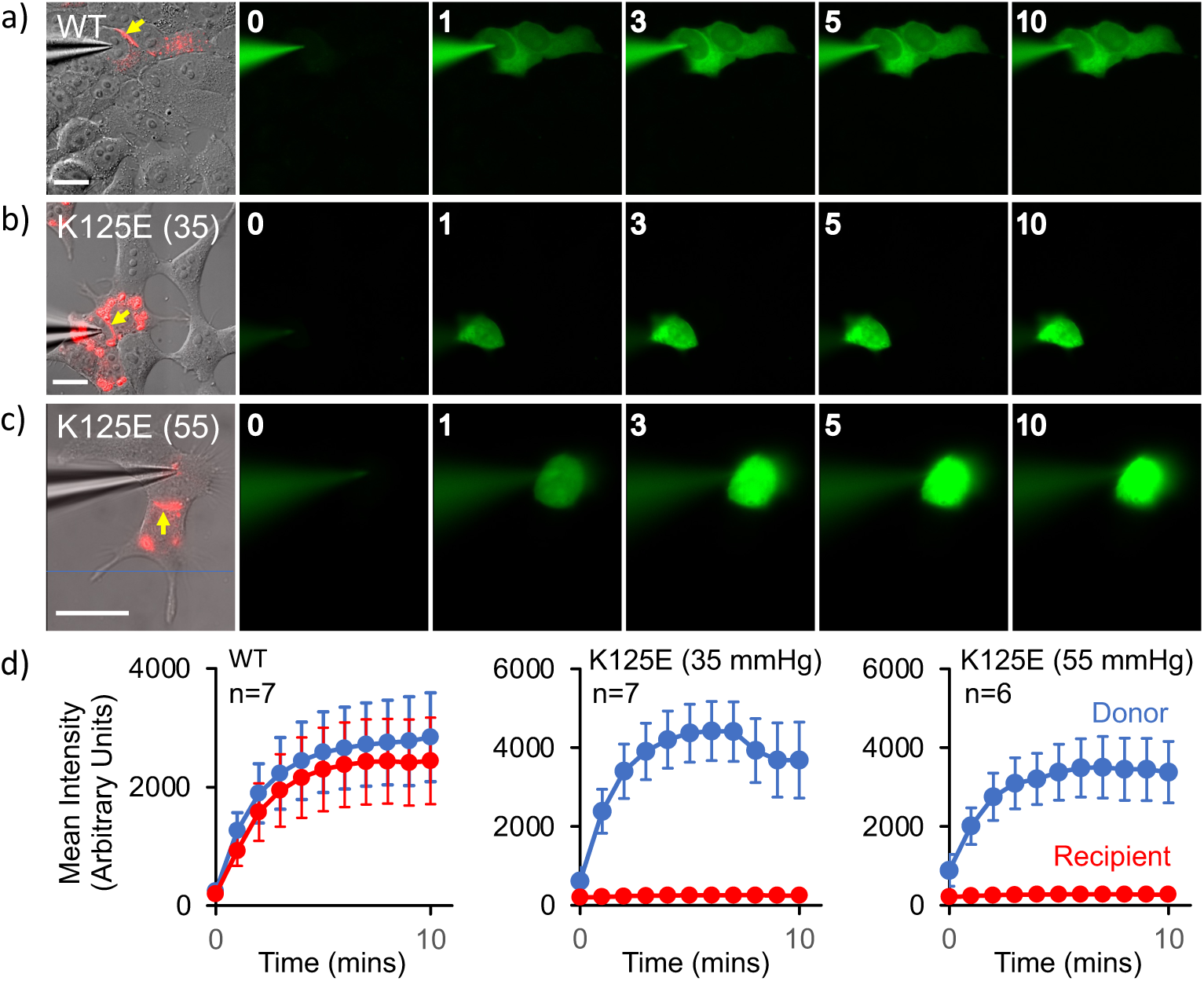
Cx26^K125E^ gap junctions are constitutively closed at a PCO_2_ of 35 mmHg. **a-c)** Montages each showing bright field DIC image of HeLa cells with mCherry fluorescence corresponding to the Cx26^K125E^-mCherry fusion superimposed (leftmost image) and the permeation of NBDG from the recorded cell to coupled cells. Yellow arrow indicates the presence of a gap junction between the cells; scale bars, 20 µm. The numbers are the time in minutes following the establishment of the whole cell recording. In Cx26^WT^ expressing cells (a), dye rapidly permeates into the coupled cell. For Cx26^K125E^ expressing cells, no dye permeates into the neighbouring cell even after 10 minutes of recording at either 35 mmHg (b) or 55 mmHg (c) PCO_2_ despite the presence of morphological gap junctions. **d)** Quantification of fluorescence intensity in the recorded cell (donor) and the potentially coupled cell (recipient) for both Cx26^WT^ and Cx26^K125E^ (7 pairs of cells recorded for WT and K125E at 35 mmHg and 6 pairs of cells for K125E at 55 mmHg, data presented as mean ± SEM). While dye permeation to the recipient cell follows the entry of dye into the donor for Cx26^WT^, no dye permeates to the acceptor cell for Cx26^K125E^. Note that the fluorescence intensity in the donor cell for Cx26^K125E^ at both levels of PCO_2_ is higher than for Cx26^WT^ at 35 mmHg, presumably because the dye remains trapped in the donor cell rather than diffusing to the recipient cell.

### The K125E mutation biases the conformational equilibrium to the N_Const_ structure

Given that the K125E mutant resulted in constitutively closed channels and the K125R mutant in channels that do not close in response to CO_2_, we set out to solve the respective structures. With respect to the wild-type and K125R constructs, purification of the K125E protein resulted in higher yields, consistent with a more stable protein. For both proteins cryo-EM data were collected from protein solubilised in DDM and vitrified in CO_2_/HCO_3_^-^ buffers corresponding to a PCO_2_ of 90mmHg with the pH maintained at pH 7.4 as was done previously (Brotherton et al., 2022). Refinement with D6 symmetry imposed resulted in maps with nominal resolutions of 2.2Å and 2.1Å respectively as defined by gold standard Fourier Shell Correlations (Rosenthal & Henderson, 2003; Scheres, 2012) (Table 1, Figure 5 – figure supplements 1 and 2). Superposition of the two maps showed there was a small but distinct change in the position of the cytoplasmic portion of TM2 between the two D6 averaged maps (Figure 5a, 5b, Figure 5 – video 1). Of the two, the K125R map looked much more similar to the equivalent map from the wild-type protein purified in the same way in DDM (PDB ID 7QEQ) and vitrified at the same PCO_2_ (Figure 5c). Further classification focussed on the cytoplasmic region of one hemichannel of the GJCs provided further evidence of a distinct difference in the conformations of the proteins. For the K125E data set the most populated class (43% of the particles) had a conformation similar to the LMNG-N_Const_ (Figure 5 – figure supplement 1). In contrast only 10% of the data for the K125R belonged to this class (Figure 5 – figure supplement 3).

**Figure 5:**
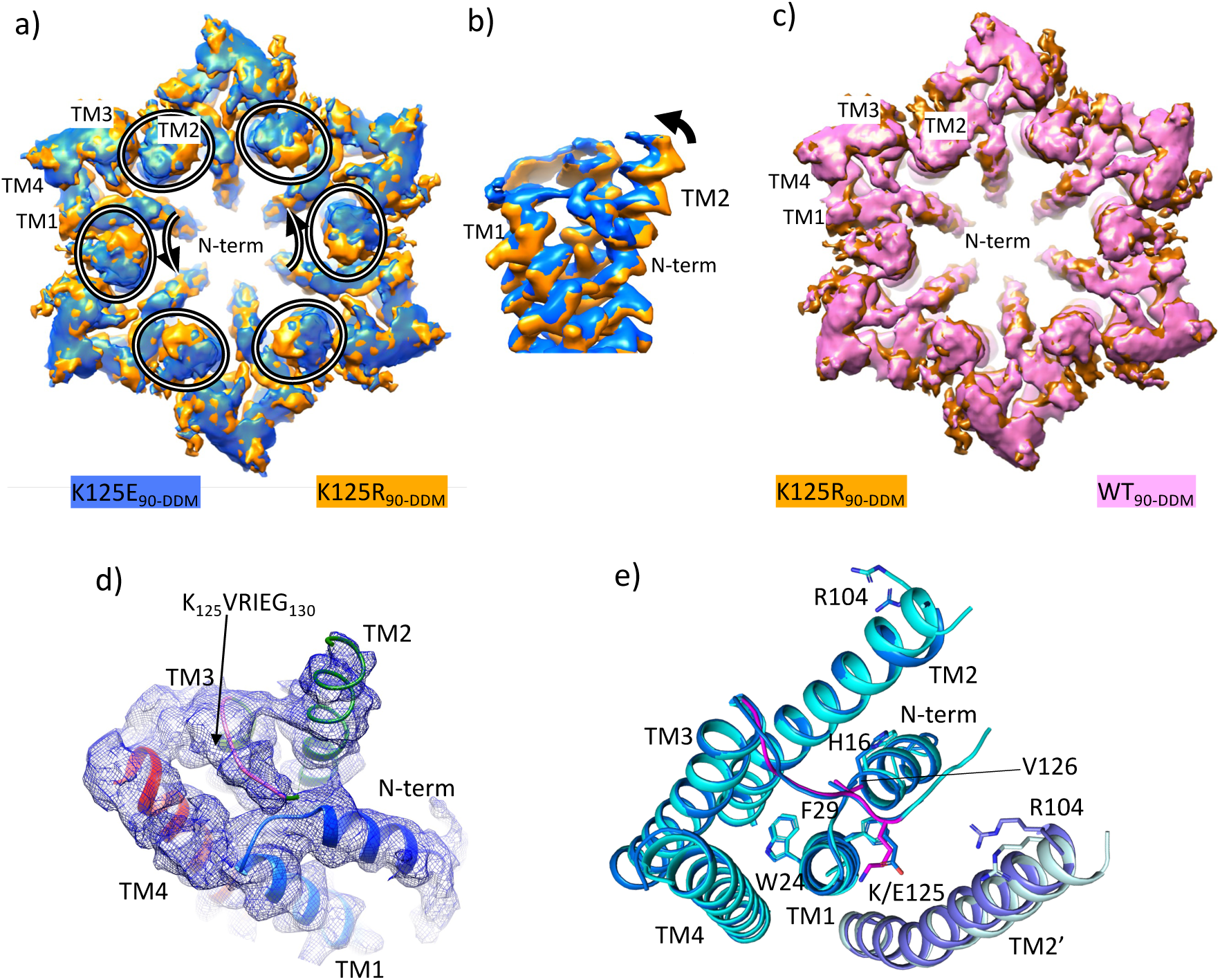
Density associated with the K125E mutant. **a)** Superposition of density for K125E_90_ D6 averaged map (blue) on the density for the K125R_90_ D6 averaged maps (orange). The ovals show the position of TM2 from each subunit and the arrows show the direction of the difference between TM2 in the two structures. **b)** As (a) but focussed on TM2 in a view approximately perpendicular to the membrane. **c)** Superposition of density for WT_90_ Cx26 (PDB ID 7QEQ; pink) D6 averaged maps on the density for R125E_90_ (orange). **d)** Density associated with one subunit of the K125E_90_ structure (unsharpened map). The structure has been coloured as in Figure 1d. **e)** Superposition of K125E_90_ structure (light blue) on the structure of LMNG-N_Const_ (cyan) showing the similarity between the two structures.

Reconstructions from the most predominant class from the K125E data set (K125E_90_) have a nominal resolution of 2.5Å (Table 2, Figure 5 – figure supplement 1). The KVRIEG motif and the N-terminus are reasonably well-defined in the density as seen for the LMNG-N_Const_ structure (Figure 5d) though again, the first three residues are difficult to position. Overall, the conformation is very similar to LMNG-N_Const_ with an RMSD of 0.35Å for 195 out of 199 matched Cα pairs (Figure 5e). In contrast the RMSD compared to the LMNG-N_Flex_ conformation is 2Å across 180 Cα pairs.

The difference of the K125E maps from the equivalent maps from the WT and K125R proteins indicated a clear bias towards a more closed conformation of the protein in the CO_2_/HCO_3_^-^ buffers. To understand the contribution of the K125E mutant, independently of any effect of CO_2_ we also reanalysed data collected from both WT and K125E protein vitrified in HEPES buffer at pH 7.4. While the resolution of the maps was much lower for these reconstructions (Table 1, Figure 5 – figure supplement 4) and better for the K125E mutant (4.2Å) than the WT protein (4.9Å) superposition of the respective maps again showed a movement of the cytoplasmic portion of TM2 together with a change in the N-terminus (Figure 5 – figure supplement 5 and Figure 5 – video 1). Thus, it appears that the K125E mutant is sufficient by itself to bias the conformation in the absence of CO_2_. However, CO_2_ may have other effects on the protein to give the higher resolution and more defined conformation seen in the CO_2_/HCO_3_^-^ buffers.

## Discussion

Connexins open and close to various stimuli. Despite a number of recent high resolution structural studies of connexins, the structural basis for pore closure or opening in response to a stimulus is still poorly understood. Cx26 is affected by PCO_2_. We have previously shown that there is a correlation in the closure of the Cx26 GJC with the level of PCO_2_ and also with the position of TM2 (Brotherton et al., 2022). Our improved data show that, not only does pore closure and the position of the N-terminal helix correlate with TM2, but also with the conformation of TM1 and importantly, the KVRIEG motif on the regulatory cytoplasmic loop.

The conformation of the KVRIEG motif and the cytoplasmic loop is interesting. In the original low resolution crystal structure of Cx26 (Cx26-xtal)(Maeda et al., 2009), the KVRIEG motif is modelled as part of TM3. However, in most other structures of connexin GJCs, the equivalent residues are not observed in the density, and it appears there is a breakdown in helical conformation at this point. In the N_Const_ structures, the KVRIEG motif forms an extended conformation mimicking the conformations seen in AlphaFold structures. In fact, the conformation of the complete loop that we observe in the subunit focussed classification, is similar to that seen in AlphaFold structures (Varadi et al., 2022). Essentially the KVRIEG motif can be described as a break in TM3 as the helix extends at either side of this extended motif. Although the low resolution of the density of the complete loop prevents the accurate positioning of the residues, the combination of our structure and the predicted models paves the way for further studies of the importance of particular interactions in the loop.

From a comparison of our structures with those of other connexins it is apparent that the position of TM2 seen in the N_Const_ structures is an outlier, with a rotation of the cytoplasmic region of the helix compared to other structures (Figure 6a). With respect to the N-terminus the various reported structures can be considered in three main categories: those with the N-terminus folded into the channel pore (N-down) (Flores et al., 2020; H. J. Lee et al., 2023; S. N. Lee et al., 2023; Myers et al., 2018); those where the N-terminus is lifted (N-up) (H. J. Lee et al.; Lee et al., 2020; Qi, Acosta Gutierrez, et al., 2023) and those where the N-terminus is flexible (N-flex) (Bennett et al., 2016; Brotherton et al., 2022; Khan et al., 2020; S. N. Lee et al., 2023; Qi, Lavriha, et al., 2023). The conformation of the Cx26-N_Const_ structure most resembles the structures with N-down as these have similar conformations of TM1 (Figure 6). However, whereas the N-terminus in these latter structures folds flat against the pore of the channel to give an open aperture, in the Cx26-N_Const_ structure, it would be prevented from doing so by the position of TM2 and so is forced outwards to form a much more constricted aperture. The conformation of TM1 in the Cx26-N_Flex_ structures, which we consider to be open, is more similar to structures of other connexins in the N-flex or N-up conformations. For Cx36 and Cx43 these conformations have been suggested to represent the closed conformation (H. J. Lee et al., 2023; Qi, Acosta Gutierrez, et al., 2023), though for Cx32 this has been described as an open conformation (Qi, Lavriha, et al., 2023). The apparent outlier amongst the Cx26 structures is the Cx26-xtal structure where the N-terminus points into the pore, but the overall conformation is more similar to the Cx26-N_Flex_ conformation. However, relative to this latter structure the helix is pulled back (Figure 6b) so is more open than seen in the Cx26-N_Const_ structure.

**Figure 6:**
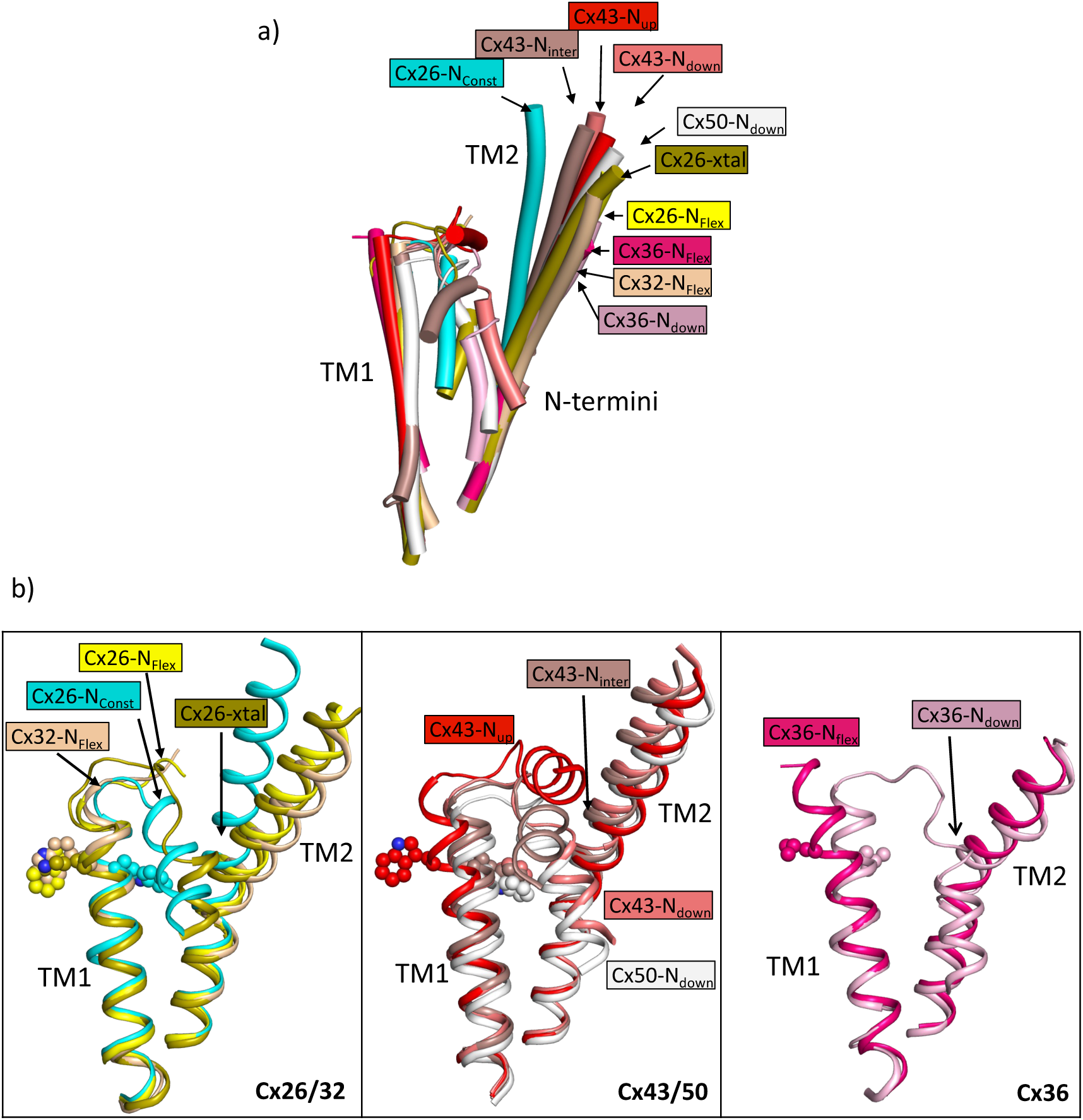
Comparison of the two structures derived from the LMNG classification with other structures of connexins. **a)** Superposition of a single subunit from the LMNG-N_Const_ (cyan) and LMNG-N_Flex_ (yellow) structures on: Cx26 crystal structure (chartreuse, PDB ID 2ZW3); Cx32 (wheat, 7zxm) Cx50 (white, 7JJP); Cx43 in up (red, 7XQF), intermediate (chocolate, 7XQI) and down (salmon, 7F94) conformations; Cx36 in down (pink, 7XNH) and flexible (raspberry, 7XKT) conformations. The structures were superposed based on all chains of the hexamer. For clarity, only TM1, TM2 and the N-terminal helices are shown for each structure. **b)** As (a) for the beta connexins Cx26 and Cx32 (left), alpha connexins Cx43 and Cx50 (middle) and the gamma connexin, Cx36 (right) structures separately. Trp24 in each of the Cx26 and Cx43 structures has been depicted with a sphere representation. The isoleucine in the corresponding position is shown for Cx36. The sequence identities for common residues to Cx26 are 63% for Cx32, 49% for Cx50, 43% for Cx43 and 35% for Cx36.

Previously, when examining our previous Cx26 maps from DDM solubilised protein we noted a lipid-like molecule in the pore of the protein and we questioned whether this would be detergent or lipid and whether it would interfere with the position of the N-terminal helix (Brotherton et al., 2022). The fact that similar density remains when the protein is solubilised with LMNG rather than DDM strongly suggests that this molecule cannot be detergent and is more likely to be a lipid. Though the presence of the molecule may still be an artefact of either the solubilisation process or heterologous expression in insect cells, it is interesting to note that a lipid-like molecule in this position appears to be a constant feature of all the high resolution cryo-EM maps associated with the structures of connexins where the N-terminus is not in the N-down position. This includes both gap junction channels (H. J. Lee et al., 2023; S. N. Lee et al., 2023; Qi, Acosta Gutierrez, et al., 2023; Qi, Lavriha, et al., 2023) and hemichannels (Lee et al., 2020; Qi, Lavriha, et al., 2023) and is irrespective of the solubilisation method or whether the structure has been solved in the presence of detergent or nanodiscs. It seems that the lipid is only displaced when the N-terminus adopts the pore lining position. As this position is not possible in the Cx26-N_Const_ structures due to the presence of TM2, the lipid remains.

It must be asked whether the N_Const_ structures that we observe represent the closed state. It is difficult to model the first three residues of the N-terminus unambiguously, presumably due to the breakdown in 6-fold symmetry at this point, but with minor changes to the side chains, the centre of the pore could be closed. The presence of the lipid would seal any other apertures between the neighbouring N-termini. Rather surprisingly given that we can isolate a symmetrical conformation of the protein using C6 symmetry, reconstructions of the full dodecamer following the focussed classifications of the single subunit do not show an influence of the conformation of that subunit on its neighbours. While this is consistent with structural studies on Cx43 gap junctions (H. J. Lee et al., 2023) the stochastic nature of the subunit conformations contrasts with studies of Cx26 hemichannels in cell membranes where significant cooperativity has been observed (de Wolf et al., 2017).

There is a clear correlation between the results of the dye transfer assays and the structural results, supporting the interpretation that the N_Const_ conformation is more like the closed state. The effect on the structure of Cx26 in changing K125 to a glutamate both in CO_2_/HCO_3_^-^ and in HEPES buffer is consistent with the hypothesis that the CO_2_ mediated closure of the gap junctions is caused by the carbamylation of K125. It would also suggest that even under high PCO_2_ conditions the WT protein is not completely carbamylated during vitrification. Given that the carbamylation reaction is highly labile, this is perhaps not surprising. The hypothesis has been that the negative charge introduced onto the lysine side-chain would enable it to form a salt bridge with Arg104 from the neighbouring subunit (Meigh et al., 2013). Consistent with this, mutation of Arg104 to alanine abolishes CO_2_ sensitivity in both GJCs (Nijjar et al., 2021) and hemichannels (Meigh et al., 2013), similar to the mutation of Lys125. In models from AlphaFold, which differ in conformation from the structure upon which the hypothesis was based, it is also notable that the lysine is located next to the arginine (Figure 3a). In the N_Const_ structures the position of the Lys/Glu125 main chain is near to Arg104, even though TM2 moves further from TM3 in moving from the N_Flex_ to the N_Const_ conformations. Though there is no clear interaction between the two residues, with minor adjustments of the side-chains, the residues could be made to interact and the absence of a definite interaction may be due to distortions during the vitrification process or radiation damage to which acidic residues are more prone. Glutamate is also much shorter than the carbamylated lysine, so that while the charges would be equivalent, the two residues might not be able to make the same specific interactions. Our results here show that the negative charge at residue 125 is important for altering conformation. For hemichannels, a direct interaction between the two residues (Meigh et al., 2015) is strongly supported by a mutant in which both R104 and K125 are replaced by cysteines, allowing a potential cross-link through a disulphide bond. This mutant responds to a change in redox potential in a similar way to which the wild type protein responds to CO_2_ (Meigh et al., 2015). However, although both Lys125 and Arg104 are necessary for CO_2_ dependent gap junction closure, attributing the induction of the conformational change in the gap junction to a single salt-bridge between Lys125 and Arg104 may be an over-simplification. For example, there may be other interactions involved and it is possible that multiple carbamylation events contribute to gap junction sensitivity (Brotherton et al., 2020). Given that we observe some protein in the N_Const_ conformation even when Arg125 is mutated to arginine, our data would be consistent with a conformationally flexible protein, in which the introduction of a negative charge would stabilise one particular conformation, rather than causing a conformational change *per se* (Figure 7).

**Figure 7:**
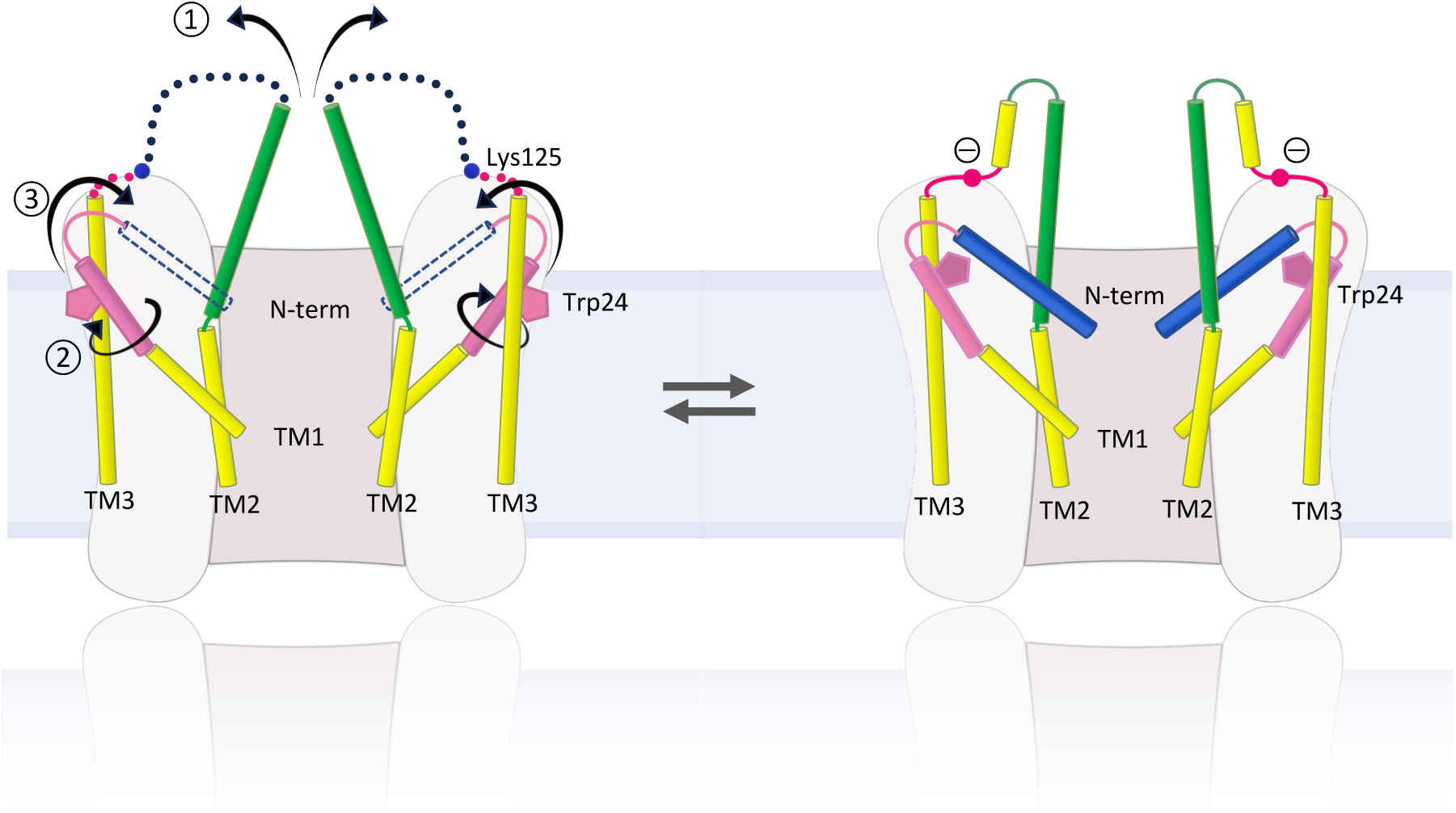
Schematic representation of conformational changes. Schematic view of the cytoplasmic region of two opposing subunits within one hemichannel of the gap junction. The open structure on the left and the constricted structure on the right are in a dynamic equilibrium. The introduction of a negative charge on Lys125 of the KVRIEG motif (magenta) pushes the equilibrium to the right. In going from one conformation to the other: ① the cytoplasmic region of TM2 (green) flexes around Phe83 and the cytoplasmic loop adopts a more defined conformation; ② the cytoplasmic region of TM1 (pink) rotates, illustrated here by the movement of Trp24; ③ the N-terminal helix (blue), which will be affected by both ① and ②, adopts a position within the pore that constricts the entrance to the channel.

Mutations in Cx26 lead to both syndromic and non-syndromic deafness (Xu & Nicholson, 2013). While these mutations have been mapped on the structure previously, the position of Ala88, mutation of which to Val causes KIDS, is interesting with regard to the new conformations of TM1. This mutation, which is at the flexion point of TM2 (Figure 2b), results in leaky hemichannels (Mhaske et al., 2013) and has been shown to prevent CO_2_ sensitivity either to the gap junctions, in closing the channels (Nijjar et al., 2021), or to the hemichannels in opening the protein (Meigh et al., 2014). In the N_const_ structures the C_β_ of Ala88 lies within 4.2Å of Trp24, which moves during the conformational change of TM1. Replacement of the alanine with the bulkier valine would hinder this conformation from being adopted and therefore, would disfavour the closed conformation. Interestingly the alanine and the tryptophan are located next to Arg143. Mutation of Arg143 to tryptophan is a very common mutation that leads to non-syndromic hearing loss (Xu & Nicholson, 2013). Overall our data, summarised in Figure 7, provide important mechanistic insight into the conformational changes behind pore closure in Cx26, which is useful for understanding how mutations in the protein can lead to disease.

## Methods

### Mutant preparation

K125R and K125E mutations of human connexin 26 were prepared using the QuikChange II mutagenesis kit (Agilent) and the following primers: K125R forward: 5’-tcgaggagatcaaaacccagagggtccgcatcg-3’, K125R reverse: 5’-cgatgcggaccctctgggttttgatctcctcga-3’, K125E forward: 5’-gagatcaaaacccaggaggtccgcatcgaa-3’, K125E reverse: 5’-ttcgatgcggacctcctgggttttgatctc-3’ (Sigma) with the wild-type human connexin 26 pFast construct used for previous studies as the template for mutagenesis (Brotherton et al., 2022). Viruses harbouring the connexin constructs were prepared and protein expressed in *Sf9* cells.

### HeLa cell culture and transfection

HeLa DH (ECACC) cells were grown in DMEM supplemented with 10% fetal bovine serum, 50 μg/mL penicillin/streptomycin and 3 mM CaCl_2_. For intercellular dye transfer experiments, cells were seeded onto coverslips in 6 well plates at a density of 2×10^4^ cells per well. After 24 hours, the cells were transiently transfected with Cx26 constructs tagged at the C-terminus with a fluorescent marker (mCherry) according to the GeneJuice Transfection Reagent protocol (Merck Millipore).

### Patch clamp recording and gap junction assay

2-Deoxy-2-[(7-nitro-2,1,3-benzoxadiazol-4-yl)amino]-D-glucose, NBDG, was included at 200 µM in the patch recording fluid, which contained: K-gluconate 130 mM; KCl 10 mM; EGTA 5 mM; CaCl_2_ 2 mM, HEPES 10 mM, pH was adjusted to 7.3 with KOH and a resulting final osmolarity of 295 mOsm. A coverslip of cells was placed in the recording chamber and superfused with a control saline (124 mM NaCl, 3 mM KCl, 2 mM CaCl_2_, 26 mM NaHCO_3_, 1.25 mM NaH_2_PO_4_, 1 mM MgSO_4_ and 10 mM D-glucose saturated with 95% O_2_/5% CO_2_, pH 7.4, PCO_2_ 35 mmHg). Cells were imaged on a Cleverscope (MCI Neuroscience) with a Photometrics Prime camera under the control of Micromanager 1.4 software. LED illumination (Cairn Research) and an image splitter (Optosplit, Cairn Research) allowed simultaneous imaging of the mCherry-tagged Cx26 subunits and the diffusion of the NBDG into and between cells. Coupled cells for intercellular dye transfer experiments were selected based on tagged Cx26 protein expression and the presence of a gap junctional plaque, easily visible as a band of mCherry fluorescence. After establishing the whole cell mode of recording, images were collected every 10 seconds.

### Protein Production, Purification and Grid Preparation

Purification of all proteins were performed as previously described (Brotherton et al., 2022), briefly described here for each protein sample.

#### K125E in HEPES buffer

##### Protein Production and purification

*Sf9* cells harbouring the Cx26 virus were harvested at 72 hours post infection at 2500 x g in a Beckmann JLA 8.1000 rotor, cell pellets were snap frozen in liquid nitrogen, and stored at – 80°C until purification. All purification steps were performed on ice, or at 4°C. Cells were thawed in hypotonic lysis buffer (10 mM sodium phosphate, 10 mM NaCl, 5 mM MgCl_2_, 1 mM DTT, pH 8.0-DNAse I, cOmplete^™^ EDTA-free Protease Inhibitor Cocktail (Roche) and AEBSF) for 30 minutes before breakage using a dounce homogeniser. Membranes were separated by ultracentrifugation for 1 hour at 4 °C, 158000 x g. After resuspending the membranes in membrane resuspension buffer (25 mM sodium phosphate, 150 mM NaCl, 5 % glycerol, 1 mM DTT, pH 8.0-DNAse I, cOmplete^™^ EDTA-free Protease Inhibitor Cocktail and AEBSF) solubilisation was carried out in membrane solubilisation buffer (10 mM sodium phosphate, 300 mM NaCl, 5 % glycerol, 1 mM DTT, 1% DDM (Glycon Biochemicals GMBH), pH 8.0) for 3-4 hours, and insoluble material removed by a further 1 hour ultracentrifugation at 4 °C, 158000 x g. Soluble protein was batch-bound to pre-equilibrated HisPur Ni-NTA resin (Thermo Scientific) overnight and then poured into an Econo-Column for subsequent manual washing and elution steps. Resin was washed with 5x CV wash buffer (10 mM sodium phosphate, 500 mM NaCl, 10 mM histidine, 5 % glycerol, 1 mM DTT, 0.1 % DDM, pH 8.0) before eluting hCx26 with elution buffer (10 mM sodium phosphate, 500 mM NaCl, 200 mM histidine, 5 % glycerol, 1 mM DTT, 0.1 % DDM, pH 8.0). hCx26-containing fractions were dialysed (20 mM HEPES, 500 mM NaCl, 5 % glycerol, 1 mM DTT, 0.03 % DDM, pH 8.0) overnight with thrombin at (a 1:1 w/w ratio). The hCx26 was then passed through a 0.2 μm filter, concentrated using a Vivaspin concentration with 100,000 MWCO and loaded onto a Superose 6 Increase 10/300 size exclusion chromatography column (GE Healthcare Lifescience) equilibrated with the same HEPES-dialysis buffer to remove thrombin. The protein was subsequently concentrated to ∼ 3 mg/ml. The concentrated protein was then dialysed for a minimum of 3 hours prior to grid preparation against 20mM HEPES, 250mM NaCl, 2.5% Glycerol, 5mM DTT, 0.03% DDM, 1mM CaCl_2_, pH 8.0.

##### Grid preparation

Protein (3.5mg/ml) was centrifuged at 17,200 g for 5mins at 4°C. Grids (0.6/1 quantifoil AU 300) were glow discharged for 30 seconds at 30mA. Vitrification of the protein in liquid ethane at –180°C was carried out with a Vitrobot MKIV with 3 μl protein per grid at 4°C, 100 % humidity, blot force 10, 3 seconds blotting.

##### Data Collection and Processing

Data were collected using a Titan Krios G3 on a Falcon 3 detector. Data processing was performed in Relion 3 (Zivanov et al., 2018). Movies were motion corrected with MotionCor2 (Zheng et al., 2017) and the CTF parameters estimated with CTFfind-4.1 (Rohou & Grigorieff, 2015), both implemented in Relion 3. Particles were picked from selected images using the Laplacian-of-Gaussian (LoG) picker, and serial rounds of 2D classifications on binned particles were used to filter out junk and poor particles. An initial model was generated using stochastic gradient descent, and this was used for further cleaning of particles via 3D classifications. Exhaustive rounds of 3D refinement, CTF refinement and polishing were performed on unbinned particles until no further improvement of the resolution for the Coulomb shell was gained. The resolution was estimated based on the gold standard Fourier Shell Coefficient (FSC) criterion (Rosenthal & Henderson, 2003; Scheres, 2012) with a soft solvent mask. All masks for processing were prepared in Chimera (Goddard et al., 2007; Pettersen et al., 2004). All processing was carried out without imposed symmetry until the final stage, where tests with C2, C3, C6 and D6 for refinement were carried out to look for improvements in resolution.

#### WT in HEPES buffer

All methods are as above, with the following changes: the final dialysis prior to freezing was against 20 mM HEPES, 200 mM NaCl, 1 % glycerol, 1 mM DTT, 1mM CaCl_2_, 0.03 % DDM, pH 8.0. Freezing concentration was 3mg/ml WT, and data collection was carried out using a Volta phase-plate.

#### K125E in αCSF90 buffer

All methods are as for K125E in HEPES buffer, except for the following changes: Fractions eluted from the NiNTA containing hCx26 were dialysed overnight at 4 °C against 10 mM sodium phosphate, 500 mM NaCl, 5 % glycerol, 1 mM DTT, 0.03 % DDM, pH 8.0. A Superose 6 Increase 5/150 size exclusion chromatography column (GE Healthcare Lifescience) was used to remove thrombin and exchange the buffer to αCSF90 buffer (70 mM NaCl, 5 % glycerol, 1 mM DTT, 0.03 % DDM, 80 mM NaHCO_3_, 1.25 mM NaH_2_PO_4_, 3 mM KCl, 1 mM MgSO_4_, 4 mM MgCl_2_.) K125E (3.4mg/ml) was gassed with 15% CO_2_ (3 x 12 seconds) followed by centrifugation at 17,200g for 5 mins at 4°C. Grids (0.6/1 quantifoil AU 300) were glow discharged for 1min at 30mA. Vitrification of the protein in liquid ethane/propane at –180°C was carried out with a Leica GP2 automated plunge freezer with 3 μl protein per grid at 4°C, 100 % humidity, 7 seconds blotting without sensor-blot in a 15% CO_2_ atmosphere. Data were collected using a GATAN K3 detector in super-resolution mode and were processed using Relion 4.

#### K125R in αCSF90 buffer

All methods are as for K125E in αCSF90 buffer, except for the following changes: Grids (1.2/1.3 Aultrafoil Au300) were glow discharged at 30mA for 30 seconds. Vitrification of the protein in liquid ethane at –160°C was carried out with a Vitrobot with 3 μl protein per grid at 4°C, 100 % humidity, 3 seconds blotting (force 10, 1 blot, skip transfer) in a 15% CO_2_ atmosphere. Data were collected using a K3 detector in Counting bin 1 mode. Data processing was carried out in Relion 4.

#### LMNG_90_ hCx26 WT

Preparation of LMNG_90_ hCx26 WT protein was carried out as for K125E in αCSF90 buffer, with the following changes: *Sf9* cells were lysed in αCSF90 buffer (70 mM NaCl, 5 % glycerol, 1 mM DTT, 80 mM NaHCO_3_, 1.25 mM NaH_2_PO_4_, 3 mM KCl, 1 mM MgSO_4_, 4 mM MgCl_2_, pH corrected to 7.4 by addition of CO_2_) and membranes were resuspended in (110 mM NaCl, 5 % glycerol, 1 mM DTT, 80 mM NaHCO_3_, 1.25 mM NaH_2_PO_4_, 3 mM KCl, 1 mM MgSO_4_, 4 mM MgCl_2_, pH corrected to 7.4 by addition of CO_2_) and solubilised in (500 mM NaCl, 5 % glycerol, 1 mM DTT, 80 mM NaHCO_3_, 1.25 mM NaH_2_PO_4_, 3 mM KCl, 1 mM MgSO_4_, 4 mM MgCl_2_, pH corrected to 7.4 by addition of CO_2_). Samples were taken periodically to check the pH, and re-adjusted by further addition of CO_2_ when necessary to keep the pH constant. Wash buffer for NiNTA resin (500 mM NaCl, 10mM Histidine, 5 % glycerol, 1 mM DTT, 80 mM NaHCO_3_, 1.25 mM NaH_2_PO_4_, 3 mM KCl, 1 mM MgSO_4_, 4 mM MgCl_2_, pH corrected to 7.4 by addition of CO_2_) and elution buffer (500 mM NaCl, 200mM Histidine, 5 % glycerol, 1 mM DTT, 80 mM NaHCO_3_, 1.25 mM NaH_2_PO_4_, 3 mM KCl, 1 mM MgSO_4_, 4 mM MgCl_2_, pH corrected to 7.4 by addition of CO_2_) were prepared and the pH checked just prior to use, to ensure no drifting of pH before interaction with the connexin. Selected fractions eluted from NiNTA were dialysed against (500 mM NaCl, 5 % glycerol, 1 mM DTT, 80 mM NaHCO_3_, 1.25 mM NaH_2_PO_4_, 3 mM KCl, 1 mM MgSO_4_, 4 mM MgCl_2_, pH corrected to 7.4 by addition of CO_2_). The final size exclusion step was performed in αCSF90 buffer (70 mM NaCl, 5 % glycerol, 1 mM DTT, 0.03 % DDM, 80 mM NaHCO_3_, 1.25 mM NaH_2_PO_4_, 3 mM KCl, 1 mM MgSO_4_, 4 mM MgCl_2_.) without additional CO_2_. The concentrated, pooled samples were gassed to pH to 7.4 both prior to freezing as described previously (Brotherton et al., 2022). Vitrification was carried out at 3.7mg/ml on 0.6/1 Aultrafoil grids using the method described for K125R in αCSF90 buffer. Data were collected using the K3 detector in super-resolution bin 2 mode.

##### Particle subtraction and masked classification focussed on the hemichannel

Hemichannel classifications with C6 imposed symmetry was carried out as described previously (Brotherton et al., 2022). The particles from the top class were unsubtracted, and the particles were refined with C6 symmetry, using a hemichannel mask and limited angular sampling. Local Resolution estimation was carried out in Relion.

##### Particle subtraction and masked classification focussed on a single subunit

Following particle expansion with D6 symmetry and particle subtraction with a mask encompassing a single subunit, masked, fixed angle classification was carried out in Relion 4. Following unsubtraction of particles, refinement of the particle positions was carried out as above, with a hemichannel mask.

### Model building and Refinement

Model building was carried out in Coot (Emsley & Cowtan, 2004) with real space refinement in Phenix (Emsley & Cowtan, 2004; Liebschner et al., 2019) using maps that had been sharpened using model free local sharpening in Phenix. For the LMNG_90_ hemichannel-based classification two maps were selected for refinement. The first of these (LMNG-N_Const_) was chosen because the density of the cytoplasmic region was the most defined. The second (LMNG-N_Flex_) was chosen as the highest resolution map with TM2 in the most diverse position. A similar selection was made for the single subunit-based classification. In building the cytoplasmic region of the protein reference was made to both ModelAngelo (Jamali et al., 2023) and AlphaFold (Jumper & Hassabis, 2022). AlphaFold structures were created with Colabfold (Mirdita et al., 2022) or downloaded from the EBI (Varadi et al., 2022).

### Structural Analysis

All structural images shown in this paper were generated in PyMol (Delano, 2002) or Chimera (Goddard et al., 2007; Pettersen et al., 2004). Superpositions were carried out in Chimera such that only matching C_a_ pairs within 2Å after superposition were included in the matrix calculation.

## Acknowledgements

We thank the Leverhulme Trust (RPG-2015-090) and MRC (MR/P010393/1) for support. We acknowledge the Midlands Regional Cryo-EM Facility, hosted at the Warwick Advanced Bioimaging Research Technology Platform, for use of the JEOL 2100Plus, and the Midlands Regional Cryo-EM Facility, hosted at Leicester Institute of Structural and Chemical Biology for use of the FEI Titan Krios G3 and associated facilities, supported by MRC award reference MC_PC_17136. We thank Dr TJ Ragan for help with cryo-EM. We are grateful to the technical support in the School of Life Sciences, University of Warwick.

## Author Contributions

The project was initiated and supervised by ND and AC. Cloning, expression, purification and grid preparation were carried out by DB. Data collection was performed by DB and CS. Data processing was done by DB and AC. DB and AC refined and analysed the structures. SN carried out the dye transfer assays. AC, DB and ND wrote the manuscript.

## Data Availability

Cryo-EM density maps have been deposited in the Electron Microscopy Data Bank (EMDB) under accession numbers EMD-18290 (K125E_90_), EMD-18295 (K125R_90_), EMD-18296 (K125E_HEPES_), EMD-18297 (WT_HEPES_), EMD-18291 (LMNG-N_Const_), EMD-18292 (LMNG-N_Flex_), EMD-18293 (LMNG-N_Const-mon_), EMD-18294 (LMNG-N_Flex-mon_). Structure models have been deposited in the RCSB Protein Data Bank under accession numbers 8Q9Z, 8QA1, 8QA0, 8QA1, 8QA3 as listed in Table 2.

## Supplementary Material

**Figure 1 – supplementary 1:**
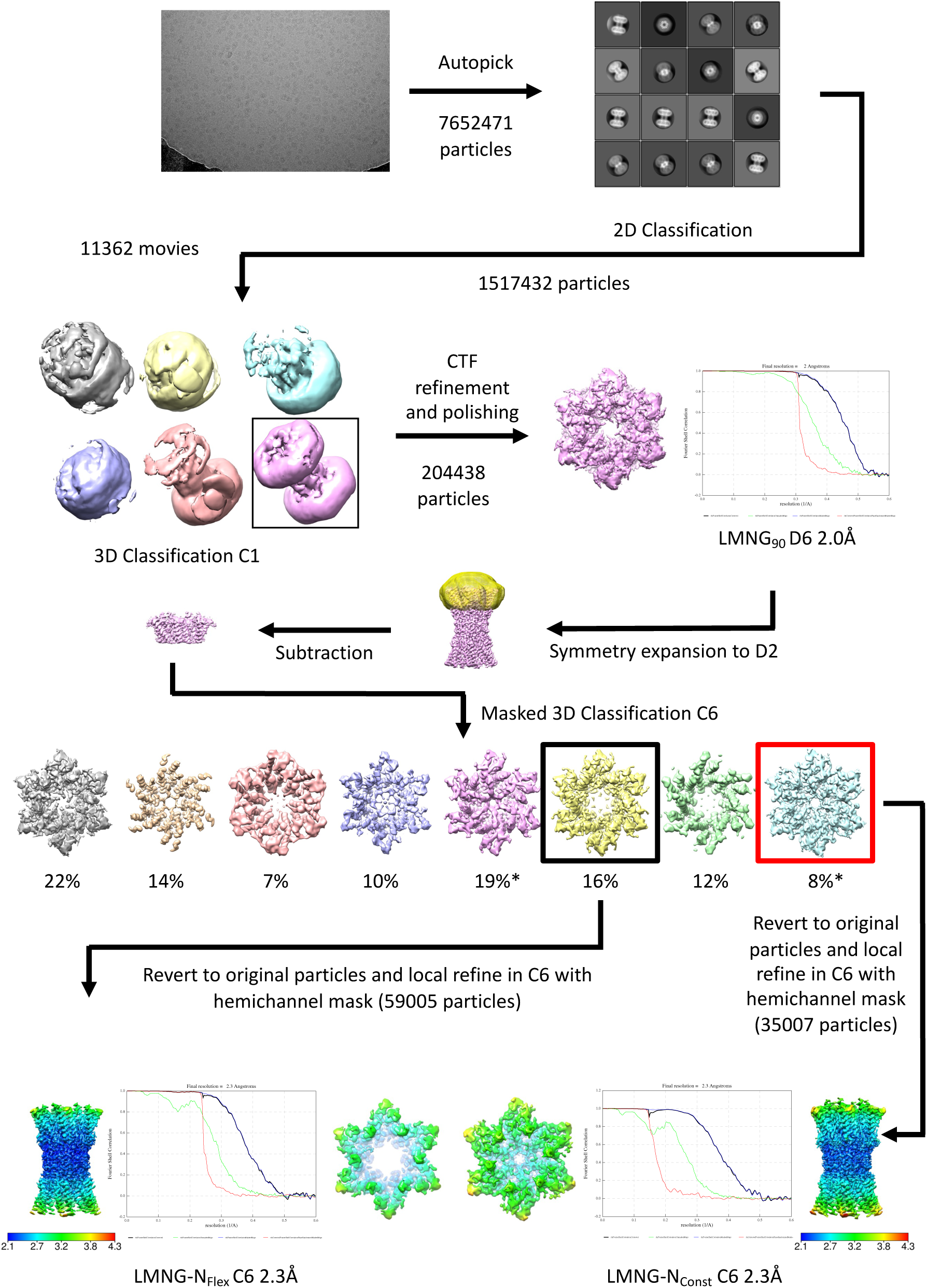
Workflow for initial processing of cryo-EM data for WT sample, solubilised in LMNG in CO_2_/HCO3^-^ buffer. The star denotes the classifications with the appearance of the NConst conformation that refine to a resolution greater than 4Å. The maps in the lower panel are coloured according to resolution as estimated in Relion 4.

**Figure 1 – supplementary 2:**
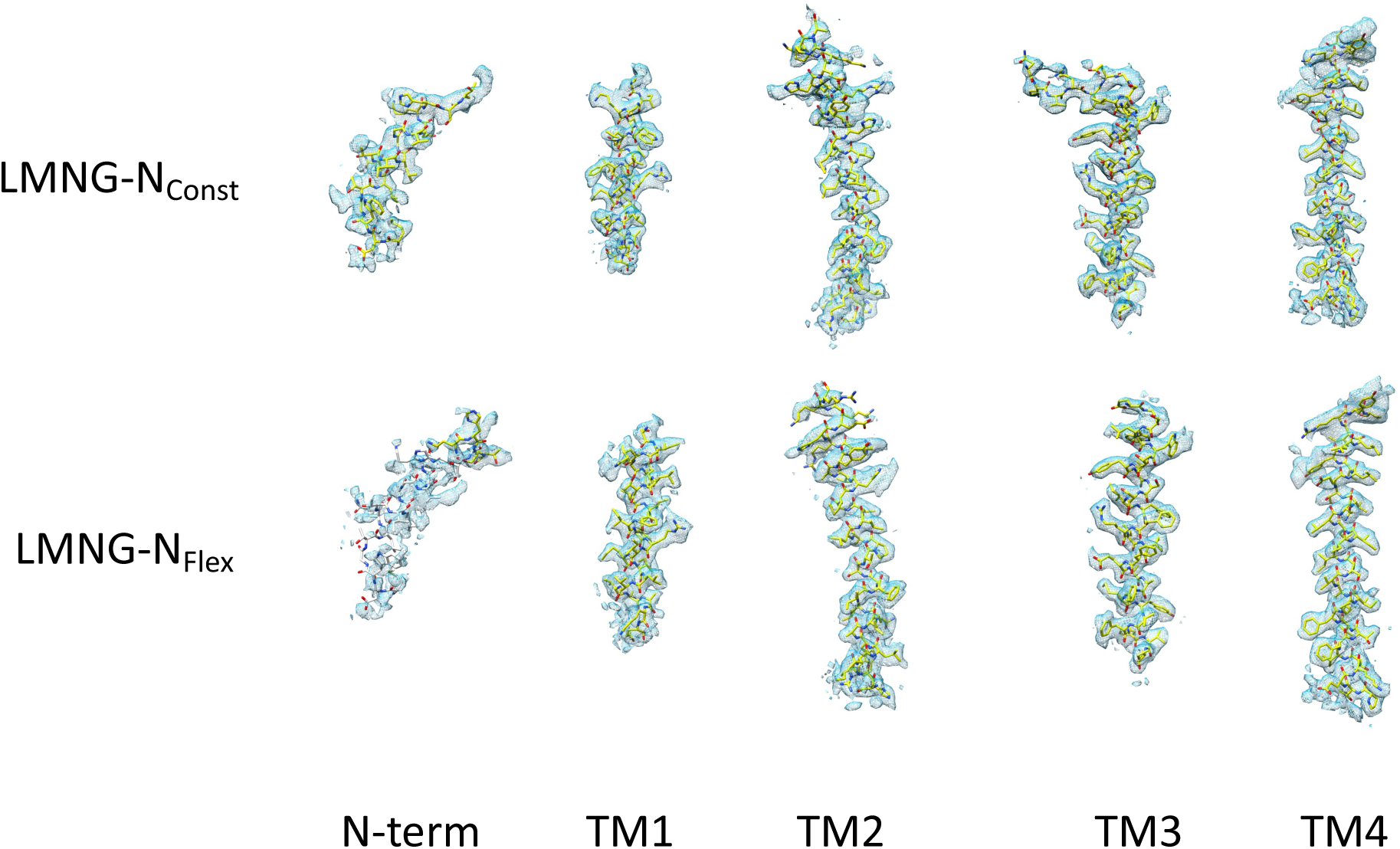
Density for the transmembrane helices and the N-terminal helix associated with each the LMNG-N_Const_ and LMNG-N_Flex_ structures. The residues with white carbon atoms are not included in the final structure.

**Figure 1 – supplementary 3:**
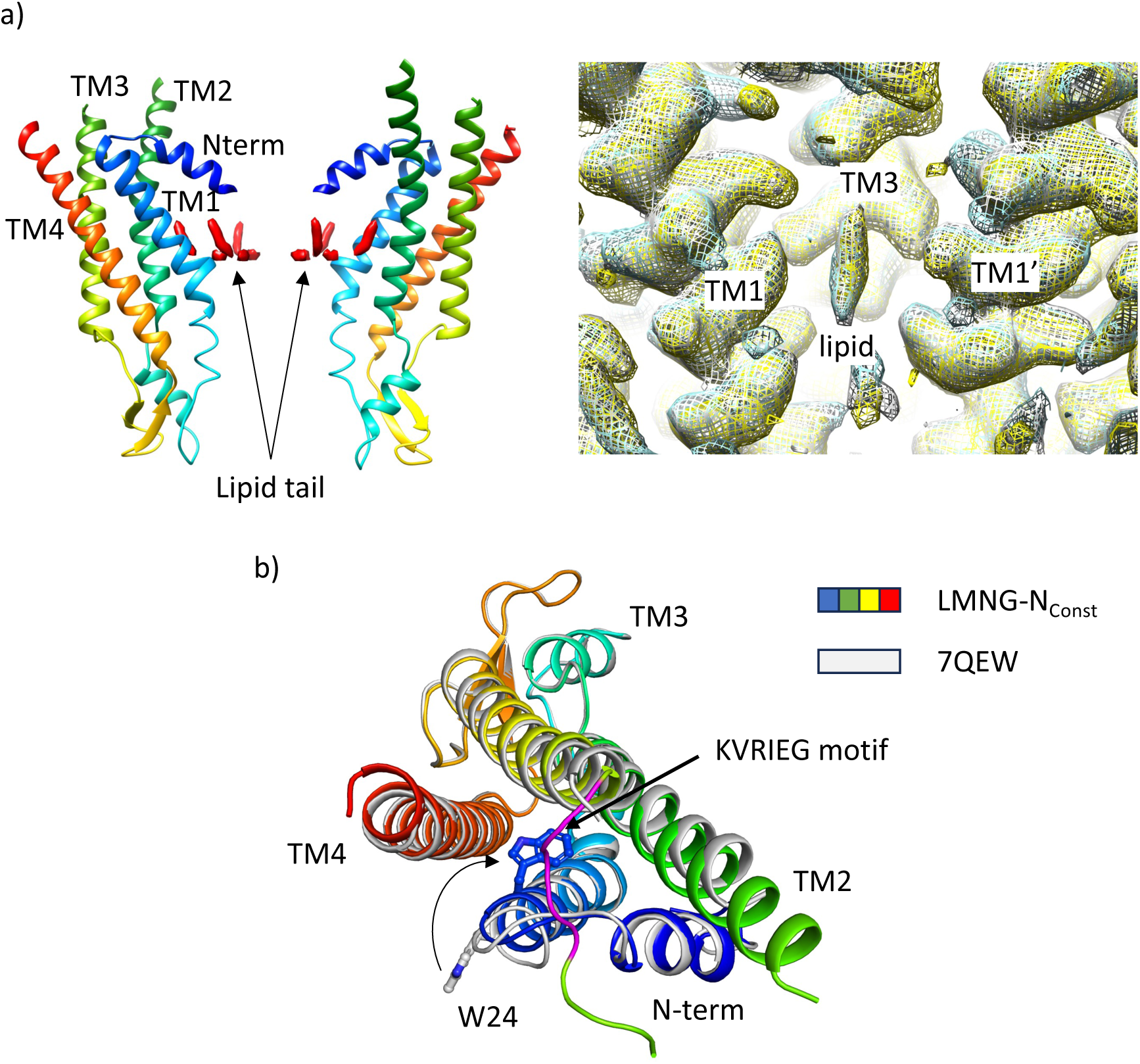
Comparison of structures derived from LMNG solubilised protein with the structure derived from the DDM solubilised protein. **a**) Density for lipid-like molecule. Left: Lipid-like density (red surface) in the pore of the protein seen in WT Cx26 solubilised in DDM (PDB 7QEQ, EMD-13937). Right: Maps associated with the LMNG solubilised protein superposed on that of the DDM solubilised protein (LMNG-_NConst_ (cyan) and LMNG-_NFlex_ (yellow), EMD-13937 (white) showing that the density remains irrespective of which of the two detergents is used to solubilise the protein. **b)** Superposition of the LMNG-N_Const_ structure on the similar conformation of the protein derived from DDM-solubilised protein (7QEW). The conformation of TM1 differs between the two structures, as shown by the position of W24. We attribute the difference to incomplete particle separation of the DDM-derived protein during the classification procedure.

**Figure 3 – supplementary figure 1:**
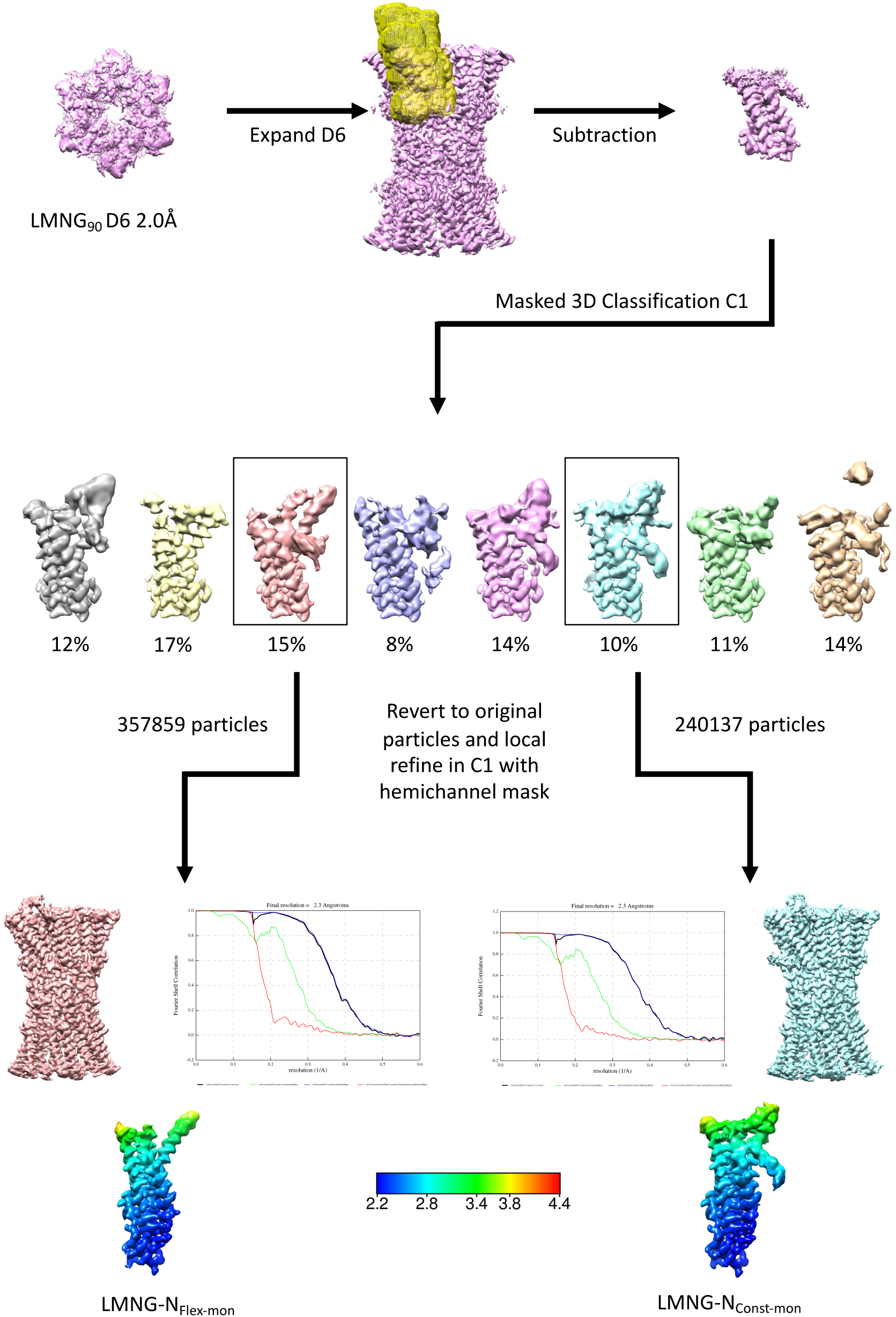
Workflow for single subunit classification for LMNG solubilised sample. The maps in the lower panel are coloured according to resolution as estimated in Relion 4.

**Figure 3 – supplementary figure 2:**
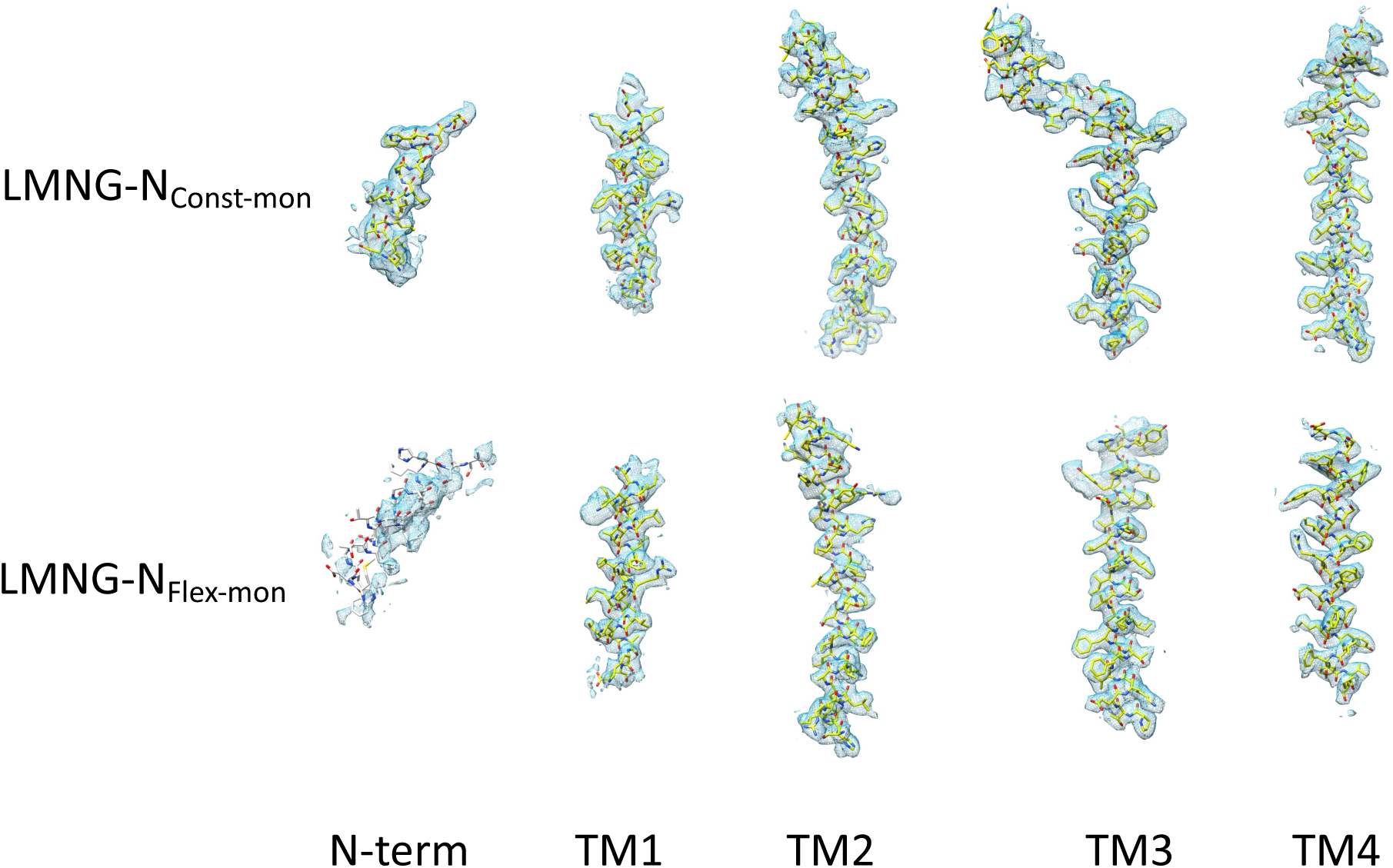
Density for the transmembrane helices and the N-terminal helix associated with each the LMNG-N_Const-mon_ and LMNG-N_Flex-mon_ structures. The residues with white carbon atoms are not included in the final structure.

**Figure 5 – supplementary figure 1:**
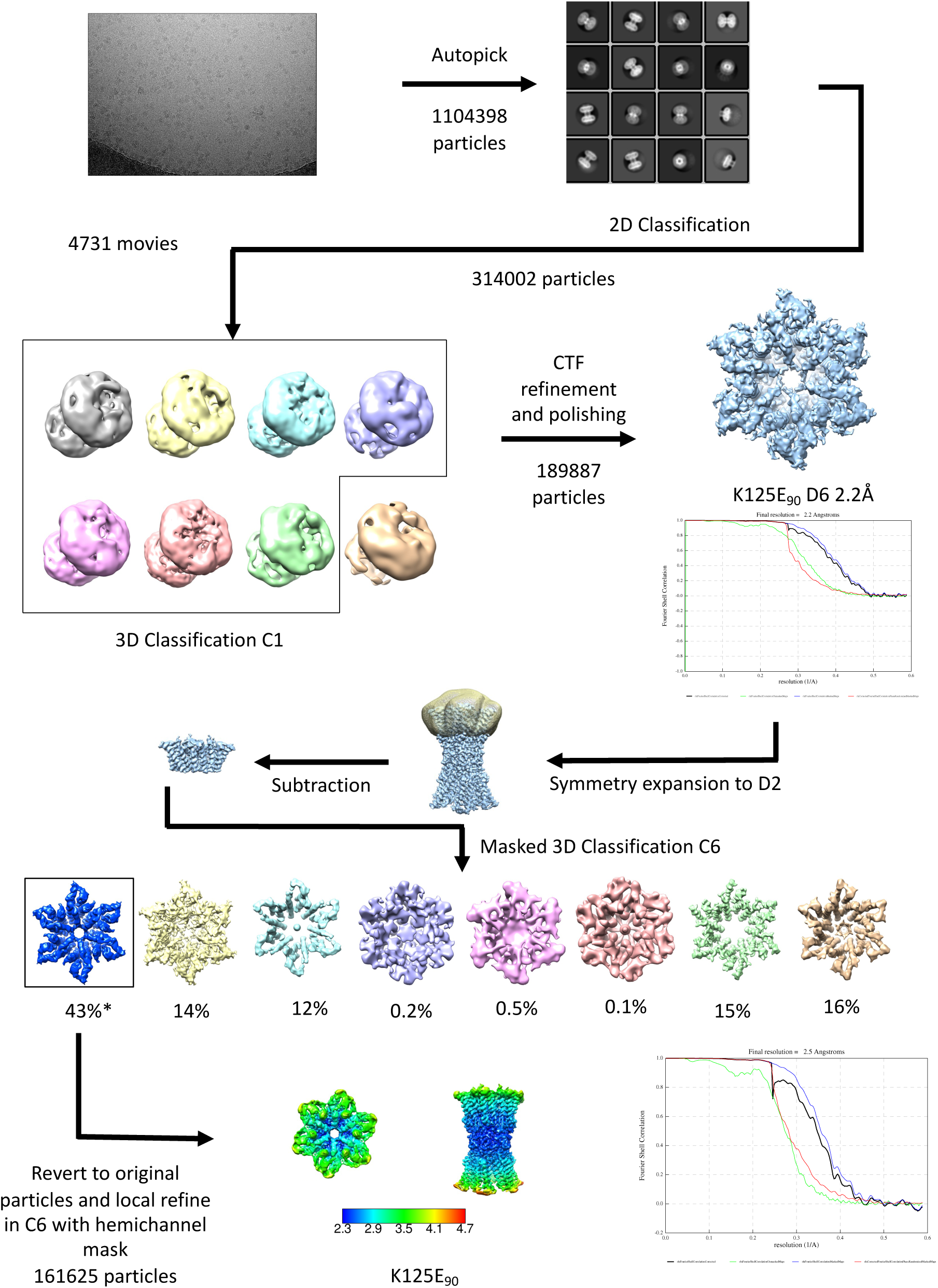
Workflow for processing of cryo-EM data for K125E sample in CO_2_/HCO3^-^ buffer. The star denotes the classifications with the appearance of the PCN conformation that refine to a resolution greater than 4Å. The maps in the lower panel are coloured according to resolution as estimated in Relion 4.

**Figure 5 – supplementary figure 2:**
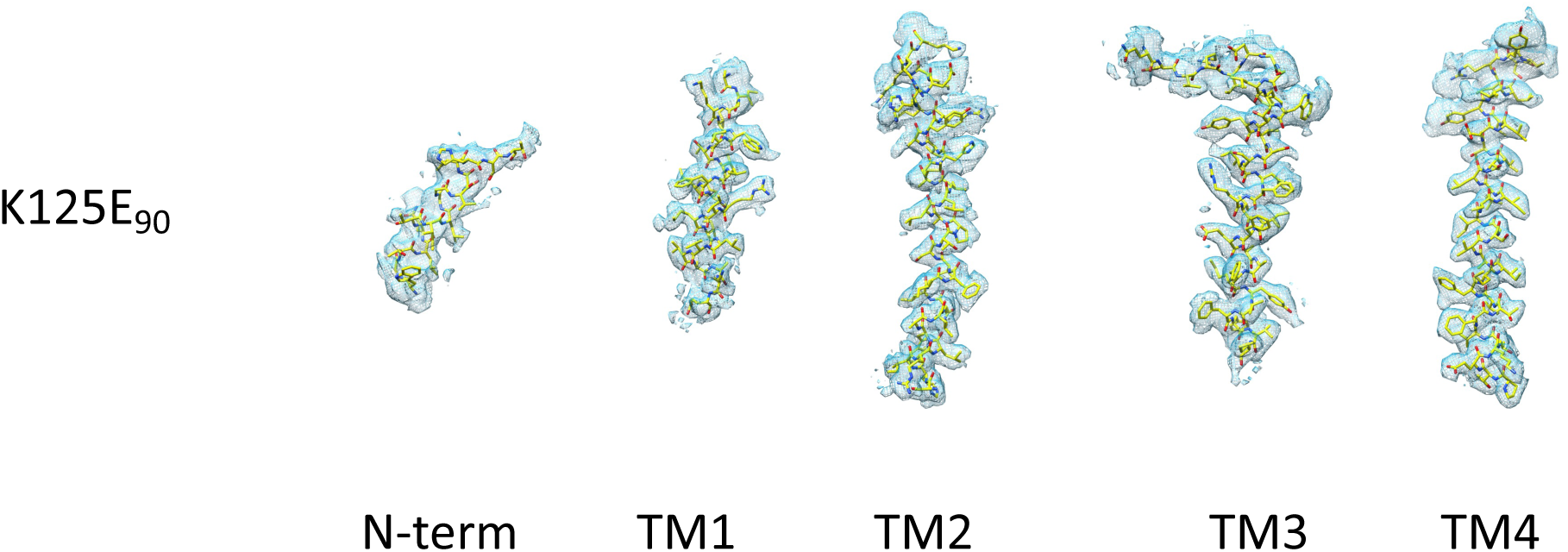
Density for the transmembrane and N-terminal helix associated with the K125E structure.

**Figure 5 – supplementary figure 3:**
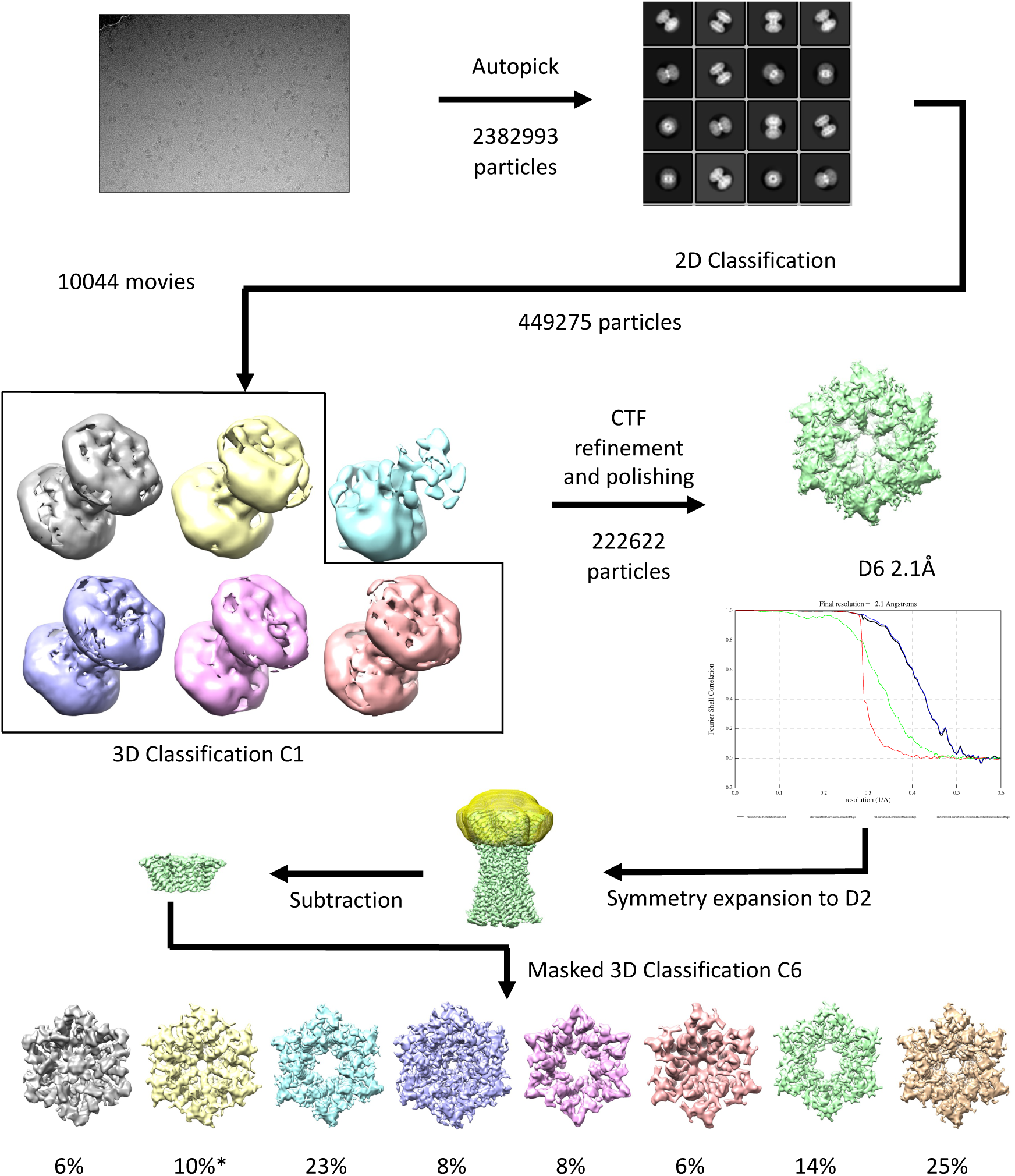
Workflow for processing of cryo-EM data for K125R sample in CO_2_/HCO3^-^ buffer. The star denotes the classifications with the appearance of the PCN conformation that refine to a resolution greater than 4Å.

**Figure 5 – supplementary figure 4:**
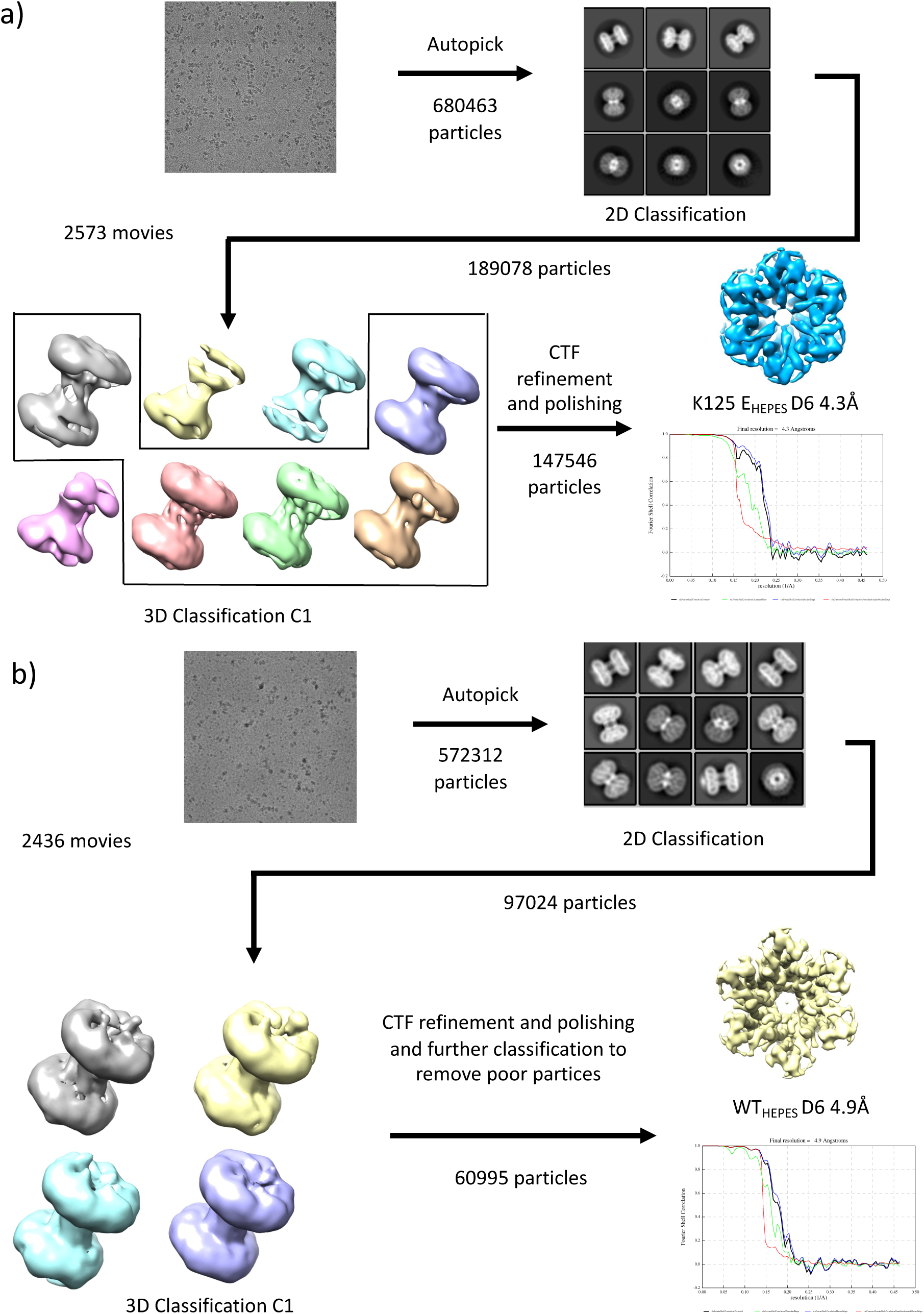
Workflows for processing of cryo-EM data for samples in HEPES buffer.

**Figure 5 – supplementary figure 5:**
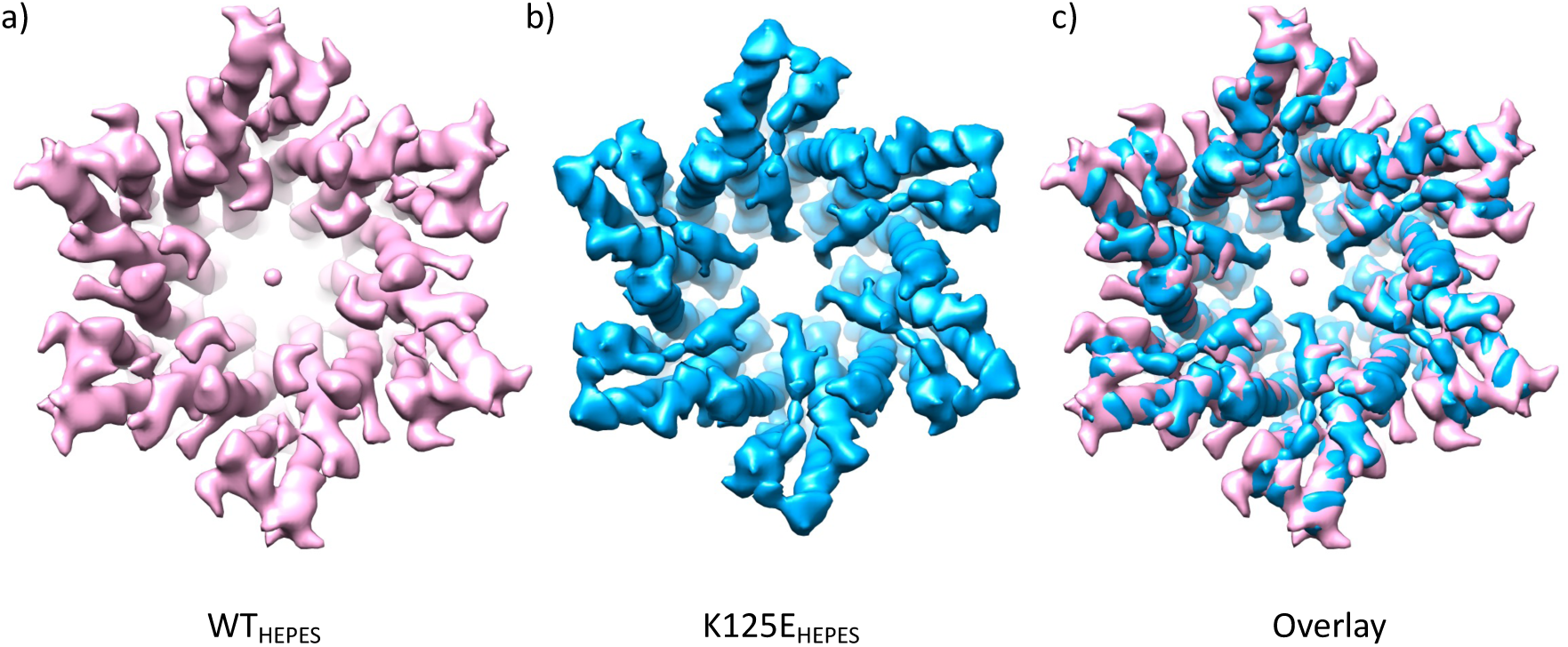
Comparison of density maps from wild type and K125E Cx26 purified in HEPES buffer at pH 7.4: **a)** WT Cx26 at 4.9Å resolution sharpened with a B-factor of –100. **b)** K125E sharpened with B-factor of –273 and low pass filtered to 5Å. **c)** Superposition of the two maps.

## Videos

**Figure 1 – video 1: Morph showing the conformational differences between reconstructions of LMNG-N_Const_ and LMNG-N_Flex_**. LMNG-N_Const_ (pale blue) LMNG-N_Flex_ (yellow).

**Figure 2 – video 1: Morph showing the conformational differences between structures of LMNG-N**_Const_ **and LMNG-N**_Flex_. As the N-terminus and KVRIEG motif are not visible in the LMNG-N_Flex_ structure these residues are not present in the morph. Residues linking the N-terminal helix to TM1 are shown in blue. Trp24 is depicted by pink sticks.

**Figure 2 – video 2: Morph showing the conformational differences between structures of LMNG-N**_Const_ **and LMNG-N**_Flex_. Focussed on TM1 with TM1, 2, 3, 4 of one subunit depicted in cyan, green, yellow and red respectively and TM2 of a second subunit in darker green. The movement of TM2 stems from the region around Phe 83 so that the position of the C_α_ atom of Lys103 near the top of TM2 of the respective structures differs by ∼8.5Å

**Figure 5 – video 1: Morph showing the conformational difference between D6 refined reconstructions of K125E and K125R.** K125E is coloured blue and K125R orange. The position of TM2 is highlighted by an oval in one of the subunits.

**Figure 5 – video 2: Morph showing the conformational differences between D6 refined reconstructions of WT and K125E in HEPES buffer.** WT (pink) K125E (blue) The position of TM2 is highlighted by an oval in one of the subunits.

## References

1. Abascal, F., & Zardoya, R. (2013). Evolutionary analyses of gap junction protein families. Biochim Biophys Acta, 1828(1), 4–14. 10.1016/j.bbamem.2012.02.007

2. Bennett, B. C., Purdy, M. D., Baker, K. A., Acharya, C., McIntire, W. E., Stevens, R. C., Zhang, Q., Harris, A. L., Abagyan, R., & Yeager, M. (2016). An electrostatic mechanism for Ca(2+)-mediated regulation of gap junction channels. Nat Commun, 7, 8770. 10.1038/ncomms9770

3. Bevans, C. G., & Harris, A. L. (1999). Regulation of connexin channels by pH. Direct action of the protonated form of taurine and other aminosulfonates. J Biol Chem, 274(6), 3711–3719. 10.1074/jbc.274.6.3711

4. Brotherton, D. H., Savva, C. G., Ragan, T. J., Dale, N., & Cameron, A. D. (2022). Conformational changes and CO(2)-induced channel gating in connexin26. Structure, 30(5), 697–706 e694. 10.1016/j.str.2022.02.010

5. Brotherton, D. H., Savva, C. G., Ragan, T. J., Linthwaite, V. L., Cann, M. J., Dale, N., & Cameron, A. D. (2020). Conformational changes and channel gating induced by CO2 binding to Connexin26. bioRxiv. 10.1101/2020.08.11.243964

6. Burendei, B., Shinozaki, R., Watanabe, M., Terada, T., Tani, K., Fujiyoshi, Y., & Oshima, A. (2020). Cryo-EM structures of undocked innexin-6 hemichannels in phospholipids. Sci Adv, 6(7), eaax3157. 10.1126/sciadv.aax3157

7. de Wolf, E., Cook, J., & Dale, N. (2017). Evolutionary adaptation of the sensitivity of connexin26 hemichannels to CO2. Proc Biol Sci, 284(1848). 10.1098/rspb.2016.2723

8. Delano, W. L. (2002). The PyMOL Molecular Graphics System. DeLano Scientific, Palo Alto, CA, USA.

9. Dospinescu, V. M., Nijjar, S., Spanos, F., Cook, J., de Wolf, E., Biscotti, M. A., Gerdol, M., & Dale, N. (2019). Structural determinants of CO2-sensitivity in the beta connexin family suggested by evolutionary analysis. Commun Biol, 2, 331. 10.1038/s42003-019-0576-2

10. Emsley, P., & Cowtan, K. (2004). Coot: model-building tools for molecular graphics. Acta Crystallogr D Biol Crystallogr, 60(Pt 12 Pt 1), 2126-2132. 10.1107/S0907444904019158

11. Flores, J. A., Haddad, B. G., Dolan, K. A., Myers, J. B., Yoshioka, C. C., Copperman, J., Zuckerman, D. M., & Reichow, S. L. (2020). Connexin-46/50 in a dynamic lipid environment resolved by CryoEM at 1.9 A. Nat Commun, 11(1), 4331. 10.1038/s41467-020-18120-5

12. Goddard, T. D., Huang, C. C., & Ferrin, T. E. (2007). Visualizing density maps with UCSF Chimera. J Struct Biol, 157(1), 281–287. 10.1016/j.jsb.2006.06.010

13. Huckstepp, R. T., Eason, R., Sachdev, A., & Dale, N. (2010). CO2-dependent opening of connexin 26 and related beta connexins. J Physiol, 588(Pt 20), 3921–3931. 10.1113/jphysiol.2010.192096

14. Jamali, K., Kall, L., Zhang, R., Brown, A., Kimanius, D., & Scheres, S. H. W. (2023). Automated model building and protein identification in cryo-EM maps. bioRxiv. 10.1101/2023.05.16.541002

15. Jumper, J., Evans, R., Pritzel, A., Green, T., Figurnov, M., Ronneberger, O., Tunyasuvunakool, K., Bates, R., Zidek, A., Potapenko, A., Bridgland, A., Meyer, C., Kohl, S. A. A., Ballard, A. J., Cowie, A., Romera-Paredes, B., Nikolov, S., Jain, R., Adler, J., … Hassabis, D. (2021). Highly accurate protein structure prediction with AlphaFold. Nature, 596(7873), 583–589. 10.1038/s41586-021-03819-2

16. Jumper, J., & Hassabis, D. (2022). Protein structure predictions to atomic accuracy with AlphaFold. Nat Methods, 19(1), 11–12. 10.1038/s41592-021-01362-6

17. Khan, A. K., Jagielnicki, M., McIntire, W. E., Purdy, M. D., Dharmarajan, V., Griffin, P. R., & Yeager, M. (2020). A Steric “Ball-and-Chain” Mechanism for pH-Mediated Regulation of Gap Junction Channels. Cell Rep, 31(3), 107482. 10.1016/j.celrep.2020.03.046

18. Lee, H. J., Cha, H. J., Jeong, H., Lee, S. N., Lee, C. W., Kim, M., Yoo, J., & Woo, J. S. (2023). Conformational changes in the human Cx43/GJA1 gap junction channel visualized using cryo-EM. Nat Commun, 14(1), 931. 10.1038/s41467-023-36593-y

19. Lee, H. J., Jeong, H., Hyun, J., Ryu, B., Park, K., Lim, H. H., Yoo, J., & Woo, J. S. (2020). Cryo-EM structure of human Cx31.3/GJC3 connexin hemichannel. Sci Adv, 6(35), eaba4996. 10.1126/sciadv.aba4996

20. Lee, S. N., Cho, H. J., Jeong, H., Ryu, B., Lee, H. J., Kim, M., Yoo, J., Woo, J. S., & Lee, H. H. (2023). Cryo-EM structures of human Cx36/GJD2 neuronal gap junction channel. Nat Commun, 14(1), 1347. 10.1038/s41467-023-37040-8

21. Liebschner, D., Afonine, P. V., Baker, M. L., Bunkoczi, G., Chen, V. B., Croll, T. I., Hintze, B., Hung, L. W., Jain, S., McCoy, A. J., Moriarty, N. W., Oeffner, R. D., Poon, B. K., Prisant, M. G., Read, R. J., Richardson, J. S., Richardson, D. C., Sammito, M. D., Sobolev, O. V., … Adams, P. D. (2019). Macromolecular structure determination using X-rays, neutrons and electrons: recent developments in Phenix. Acta Crystallogr D Struct Biol, 75(Pt 10), 861–877. 10.1107/S2059798319011471

22. Lorimer, G. H. (1983). Carbon dioxide and carbamate formation: the makings of a biochemical control system. Trends in Biochemical Sciences, 8, 65–68.

23. Maeda, S., Nakagawa, S., Suga, M., Yamashita, E., Oshima, A., Fujiyoshi, Y., & Tsukihara, T. (2009). Structure of the connexin 26 gap junction channel at 3.5 A resolution. Nature, 458(7238), 597–602. 10.1038/nature07869

24. Meigh, L., Cook, D., Zhang, J., & Dale, N. (2015). Rational design of new NO and redox sensitivity into connexin26 hemichannels. Open Biol, 5(2), 140208. 10.1098/rsob.140208

25. Meigh, L., Greenhalgh, S. A., Rodgers, T. L., Cann, M. J., Roper, D. I., & Dale, N. (2013). CO(2)directly modulates connexin 26 by formation of carbamate bridges between subunits. Elife, 2, e01213. 10.7554/eLife.01213

26. Meigh, L., Hussain, N., Mulkey, D. K., & Dale, N. (2014). Connexin26 hemichannels with a mutation that causes KID syndrome in humans lack sensitivity to CO2. Elife, 3, e04249. 10.7554/eLife.04249

27. Mhaske, P. V., Levit, N. A., Li, L., Wang, H. Z., Lee, J. R., Shuja, Z., Brink, P. R., & White, T. W. (2013). The human Cx26-D50A and Cx26-A88V mutations causing keratitis-ichthyosis-deafness syndrome display increased hemichannel activity. Am J Physiol Cell Physiol, 304(12), C1150–1158. 10.1152/ajpcell.00374.2012

28. Mirdita, M., Schutze, K., Moriwaki, Y., Heo, L., Ovchinnikov, S., & Steinegger, M. (2022). ColabFold: making protein folding accessible to all. Nat Methods, 19(6), 679–682. 10.1038/s41592-022-01488-1

29. Myers, J. B., Haddad, B. G., O’Neill, S. E., Chorev, D. S., Yoshioka, C. C., Robinson, C. V., Zuckerman, D. M., & Reichow, S. L. (2018). Structure of native lens connexin 46/50 intercellular channels by cryo-EM. Nature, 564(7736), 372–377. 10.1038/s41586-018-0786-7

30. Nijjar, S., Maddison, D., Meigh, L., de Wolf, E., Rodgers, T., Cann, M. J., & Dale, N. (2021). Opposing modulation of Cx26 gap junctions and hemichannels by CO2. J Physiol, 599(1), 103–118. 10.1113/JP280747

31. Peracchia, C. (2004). Chemical gating of gap junction channels; roles of calcium, pH and calmodulin. Biochim Biophys Acta, 1662(1-2), 61–80. 10.1016/j.bbamem.2003.10.020

32. Pettersen, E. F., Goddard, T. D., Huang, C. C., Couch, G. S., Greenblatt, D. M., Meng, E. C., & Ferrin, T. E. (2004). UCSF Chimera--a visualization system for exploratory research and analysis. J Comput Chem, 25(13), 1605–1612. 10.1002/jcc.20084

33. Qi, C., Acosta Gutierrez, S., Lavriha, P., Othman, A., Lopez-Pigozzi, D., Bayraktar, E., Schuster, D., Picotti, P., Zamboni, N., Bortolozzi, M., Gervasio, F. L., & Korkhov, V. M. (2023). Structure of the connexin-43 gap junction channel in a putative closed state. Elife, 12. 10.7554/eLife.87616

34. Qi, C., Lavriha, P., Bayraktar, E., Vaithia, A., Schuster, D., Pannella, M., Sala, V., Picotti, P., Bortolozzi, M., & Korkhov, V. M. (2023). Structures of wild-type and selected CMT1X mutant connexin 32 gap junction channels and hemichannels. Sci Adv, 9(35), eadh4890. 10.1126/sciadv.adh4890

35. Rohou, A., & Grigorieff, N. (2015). CTFFIND4: Fast and accurate defocus estimation from electron micrographs. J Struct Biol, 192(2), 216–221. 10.1016/j.jsb.2015.08.008

36. Rosenthal, P. B., & Henderson, R. (2003). Optimal determination of particle orientation, absolute hand, and contrast loss in single-particle electron cryomicroscopy. J Mol Biol, 333(4), 721–745. 10.1016/j.jmb.2003.07.013

37. Scheres, S. H. (2012). RELION: implementation of a Bayesian approach to cryo-EM structure determination. J Struct Biol, 180(3), 519–530. 10.1016/j.jsb.2012.09.006

38. Stout, C., Goodenough, D. A., & Paul, D. L. (2004). Connexins: functions without junctions. Curr Opin Cell Biol, 16(5), 507–512. 10.1016/j.ceb.2004.07.014

39. Valiunas, V. (2002). Biophysical properties of connexin-45 gap junction hemichannels studied in vertebrate cells. J Gen Physiol, 119(2), 147–164. 10.1085/jgp.119.2.147

40. Varadi, M., Anyango, S., Deshpande, M., Nair, S., Natassia, C., Yordanova, G., Yuan, D., Stroe, O., Wood, G., Laydon, A., Zidek, A., Green, T., Tunyasuvunakool, K., Petersen, S., Jumper, J., Clancy, E., Green, R., Vora, A., Lutfi, M., … Velankar, S. (2022). AlphaFold Protein Structure Database: massively expanding the structural coverage of protein-sequence space with high-accuracy models. Nucleic Acids Res, 50(D1), D439–D444. 10.1093/nar/gkab1061

41. Xu, J., & Nicholson, B. J. (2013). The role of connexins in ear and skin physiology – functional insights from disease-associated mutations. Biochim Biophys Acta, 1828(1), 167–178. 10.1016/j.bbamem.2012.06.024

42. Young, K. C., & Peracchia, C. (2004). Opposite Cx32 and Cx26 voltage-gating response to CO2 reflects opposite voltage-gating polarity. J Membr Biol, 202(3), 161–170. 10.1007/s00232-004-0727-2

43. Yu, J., Bippes, C. A., Hand, G. M., Muller, D. J., & Sosinsky, G. E. (2007). Aminosulfonate modulated pH-induced conformational changes in connexin26 hemichannels. J Biol Chem, 282(12), 8895–8904. 10.1074/jbc.M609317200

44. Zheng, S. Q., Palovcak, E., Armache, J. P., Verba, K. A., Cheng, Y., & Agard, D. A. (2017). MotionCor2: anisotropic correction of beam-induced motion for improved cryo-electron microscopy. Nat Methods, 14(4), 331–332. 10.1038/nmeth.4193

45. Zivanov, J., Nakane, T., Forsberg, B. O., Kimanius, D., Hagen, W. J., Lindahl, E., & Scheres, S. H. (2018). New tools for automated high-resolution cryo-EM structure determination in RELION-3. Elife, 7. 10.7554/eLife.42166

